# The hearing aid dilemma: amplification, compression, and distortion of the neural code

**DOI:** 10.1101/2020.10.02.323626

**Authors:** Alex Armstrong, Chi Chung Lam, Shievanie Sabesan, Nicholas A. Lesica

## Abstract

Hearing aids are the only available treatment for mild-to-moderate sensorineural hearing loss, but often fail to improve perception in difficult listening conditions. To identify the reasons for this failure, we studied the underlying neural code using large-scale single-neuron recordings in gerbils, a common animal model of human hearing. We found that a hearing aid restored the sensitivity of neural responses, but failed to restore their selectivity. The low selectivity of aided responses was not a direct effect of hearing loss per se, but rather a consequence of the strategies used by hearing aids to restore sensitivity: compression, which decreases the spectral and temporal contrast of incoming sounds, and amplification, which produces high intensities that distort the neural code even with normal hearing. To improve future hearing aids, new processing strategies that avoid this tradeoff between neural sensitivity and selectivity must be developed.

## Introduction

Hearing loss is one of the most widespread and disabling chronic conditions in the world today. Approximately 500 million people worldwide are affected, making hearing loss the fourth leading cause of years lived with disability (Wilson et al., 2017) and imposing a substantial economic burden with estimated costs of more than $750 billion globally each year (World Health Organization, 2017). Hearing loss has also been linked to declines in mental health; in fact, a recent commission identified hearing loss as the leading modifiable risk factor for incident dementia (Livingston et al., 2017). As the societal impact of hearing loss continues to grow, the need for improved treatments is becoming increasingly urgent.

Hearing aids are the current treatment of choice and remain the only option for the most common forms of hearing loss that result from noise exposure and aging. But only a small fraction of people with hearing loss have hearing aids, and only a small fraction of those with hearing aids actually use them (McCormack and Fortnum, 2013; Orji et al., 2020). There are a number of reasons for this poor uptake, but one of the most important is lack of benefit in listening environments that are typical of real-world social settings. The primary problem associated with hearing impairment is loss of audibility (Humes, 1996; Humes and Dubno, 2010). As a result of cochlear damage, sensitivity thresholds are increased and low-intensity sounds can no longer be perceived. Fortunately, hearing aids are generally able to correct this problem by providing amplification. But perception often remains impaired even after audibility is restored. It is well established that hearing aids improve the perception of low-intensity sounds in quiet environments but often fail to provide benefit for high-intensity sounds in background noise (Humes et al., 1999; Larson et al., 2000).

The reasons for this residual impairment remain unclear, but one possibility is the existence of additional deficits beyond loss of audibility that impair the processing of high-intensity sounds. Many such deficits have been reported such as broadened frequency tuning (Moore, 2007) and impaired temporal processing (Henry and Heinz, 2013; Lorenzi et al., 2006). But these deficits are typically observed when comparisons between normal and impaired hearing are made at different sound intensities to control for differences in audibility. This approach confounds the effects of hearing loss with the effects of intensity; amplification to high intensities impairs auditory processing even with normal hearing (Horvath and Lesica, 2011; Studebaker et al., 1999; Wong et al., 1998). In fact, when listeners with mild-to-moderate hearing loss (typical of the vast majority of impairments) and normal hearing listeners are compared at the same high intensities, the performance of the two groups is often similar in both simple tasks such as tone-in-noise detection (Nelson, 1991) and complex tasks such as speech-in-noise perception (Ching et al., 1998; Lee and Humes, 1993; Oxenham and Kreft, 2016; Studebaker et al., 1999; Summers and Cord, 2007).

Another possibility is that the residual problems that persist after restoration of audibility are caused by the processing in the hearing aid itself. Most modern hearing aids share the same core processing algorithm known as multi-channel wide dynamic range compression (WDRC). This algorithm provides listeners with frequency-specific amplification based on measured changes in their sensitivity thresholds. It also provides compression by varying the amplification of each frequency over time based on the incoming sound intensity such that amplification decreases as the incoming sound intensity increases. This algorithm is designed to mimic the amplification and compression that normally take place within a healthy cochlea but are compromised by hearing loss. However, it ignores many other aspects of auditory processing that are also impacted by hearing loss (Lesica, 2018) and modifies the spectral and temporal properties of incoming sounds in ways that may actually be detrimental to perception (Kates, 2010; Souza, 2002).

Identifying the factors responsible for the failure of hearing aids to restore normal auditory perception through psychophysical studies has proven difficult. We approached the problem from the perspective of the neural code -- the activity patterns in central auditory brain areas that provide the link between sound and perception. Hearing loss impairs perception because it causes distortions in the information carried by the neural code about incoming sounds. The failure of current hearing aids to restore normal perception suggests that there are critical features of the neural code that remain distorted. An ideal hearing aid would correct these distortions by transforming incoming sounds such that processing of the transformed sounds by the impaired system would result in the same neural activity patterns as the processing of the original sounds by the healthy system; current hearing aids fail to achieve this ideal.

Little is known about the specific distortions in the neural code caused by hearing loss or the degree to which current hearing aids correct them. The effects of hearing loss on the neural code for complex sounds such as speech have been well characterized at the level of the auditory nerve (Young, 2012), but its impact on downstream central brain areas remains unclear as there have been few studies of single neuron responses with hearing loss and none with hearing aids. Auditory processing in humans involves many brain areas from the brainstem, which performs general feature extraction and integration, to the cortex, which performs context- and language-specific processing. While large-scale studies of single neurons in these areas in humans are not yet possible, animal models can serve as a valuable surrogate, particularly for the early stages of processing which are largely conserved across mammals and appear to be the primary source of human perceptual deficits (Humes and Dubno, 2010). Prior work has already shown that classifiers trained to identify speech phonemes based on neural activity patterns recorded from animals perform similarly to human listeners performing an analogous task (Mesgarani et al., 2008). Thus, comparisons of the neural code with and without hearing loss and a hearing aid in an animal model can provide valuable insight into which distortions in the neural code underlie the failure of hearing aids to restore normal perception.

We focused our study on mild-to-moderate sensorineural hearing loss, which reflects relatively modest cochlear damage (Liberman et al., 1986). Because peripheral processing is still highly functional with this form of hearing loss, there is potential for a hearing aid to provide substantial benefit. We chose to study the neural code in the inferior colliculus (IC), the midbrain hub of the central auditory pathway. The IC is several processing stages downstream of the cochlea and reflects the cumulative effects of brainstem processing, but its activity is still largely determined by the acoustic features of incoming sounds rather than contextual or behavioral factors. We found that most of the distortions in the neural code that are caused by hearing loss are, in fact, corrected by a hearing aid; only a loss of selectivity in neural responses that is specific to complex sounds remains. Our analysis suggests that the low selectivity of aided responses is not a fundamental deficit in auditory processing, but is instead a consequence of the strategies used by current hearing aids to restore audibility. These findings support the wide provision of simple devices to address the growing global burden of hearing loss in the short term and provide guidance for the development of improved hearing aids in the future.

## Results

To study the neural code with high spatial and temporal resolution across large populations of neurons, we made recordings using custom-designed electrodes with a total of 512 channels spanning both brain hemispheres in gerbils, a commonly used model for studies of low-frequency hearing (Figure 1A). We used these large-scale recordings to study the activity patterns of more than 5,000 neurons in the IC. To induce sloping mild-to-moderate sensorineural hearing loss, we exposed young-adult gerbils to broadband noise (118 dB SPL, 3 hours). Compared to normal hearing animals, the resulting pure-tone threshold shifts measured one month after exposure using auditory brainstem response (ABR) recordings typically ranged from 20-30 dB at low frequencies to 40-50 dB at high frequencies (Figure 1B). Pure-tone threshold shifts with hearing loss were also evident in frequency response areas (FRAs) measured from multi-unit activity (MUA) recorded in the IC, which illustrate the degree to which populations of neurons were responsive to tones with different frequencies and intensities (Figure 1C).

**Figure 1:**
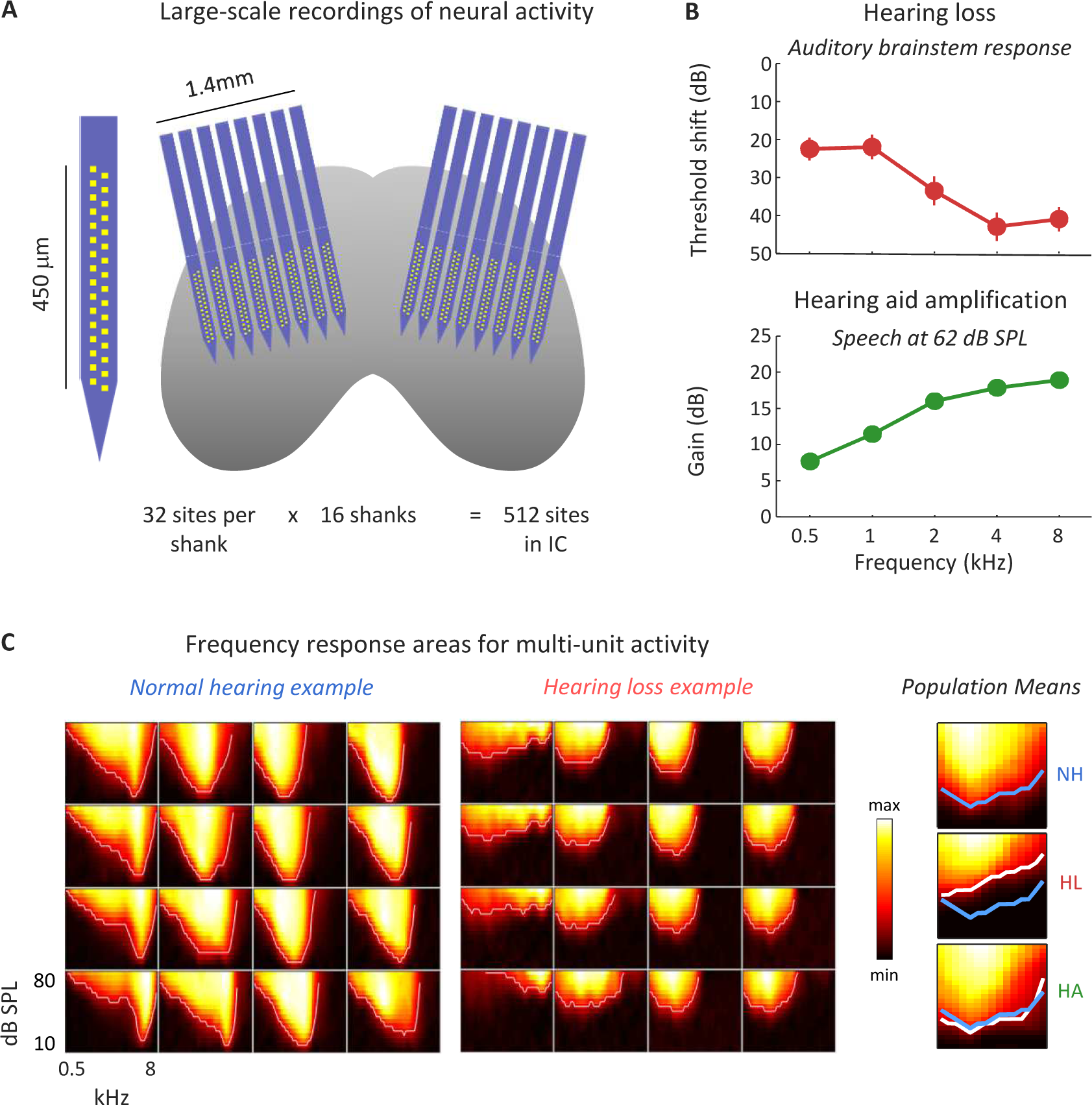
Large-scale recordings of neural activity from the inferior colliculus with normal hearing and mild-to-moderate hearing loss. (A) Schematic diagram showing the geometry of custom-designed electrode arrays in relation to the inferior colliculus. (B) Threshold shifts with hearing loss and corresponding hearing aid amplification. Top: Hearing loss as a function of frequency in noise-exposed animals (mean ± standard error, n = 20). The values shown are the ABR threshold shift relative to the mean of all animals with normal hearing (n = 15). Bottom: Hearing aid amplification as a function of frequency for speech at 62 dB SPL with gain and compression parameters fit to the average hearing loss after noise exposure. The values shown are the average across 5 minutes of continuous speech. (C) MUA recorded in the inferior colliculus during the presentation of tones. Left: The MUA FRAs for 16 channels from a normal hearing animal. Each subplot shows the average activity recorded from a single channel during the presentation of tones with different frequencies and intensities. Middle: MUA FRAs for 16 channels from an animal with hearing loss. Right: The average MUA FRAs across all channels from all animals for each hearing condition. The lines indicate the lowest intensity for each frequency at which the mean MUA was more than 3 standard deviations above the mean MUA during silence. The line for normal hearing is shown in blue on all three subplots.

For animals with hearing loss, we presented sounds both before and after processing with a multi-channel WDRC hearing aid. The amplification and compression parameters for the hearing aid were custom fit to each ear of each animal based on the measured ABR threshold shifts. The hearing aid amplified sounds in a frequency-dependent manner, with amplification for sounds at moderate intensity typically increasing from approximately 10 dB at low frequencies to approximately 20 dB at high frequencies (Figure 1B). This amplification was sufficient to restore the pure-tone IC MUA thresholds with hearing loss to normal (Figure 1C).

To begin our study of the neural code, we first presented speech to normal hearing animals at moderate intensity typical of a conversation in a quiet environment (62 dB SPL). We used a set of nonsense consonant-vowel syllables, as is common in human studies that focus on acoustic cues for speech perception rather than linguistic or cognitive factors. The set of syllables consisted of all possible combinations of 12 consonants and 4 vowels, each spoken by 8 different talkers. For individual neurons, individual instances of different syllables elicited complex response patterns (Figure 2A). For a population of neurons, the response patterns can be thought of as trajectories in a high-dimensional space in which each dimension corresponds to the activity of one neuron and each point on a trajectory indicates the activity of each neuron in the population at one point in time. To visualize these patterns, we performed dimensionality reduction via principal component analysis, which identified linear combinations of all neurons that best represented the full population. Within the space defined by the first three principal components, the responses to individual instances of different syllables followed distinct trajectories that were reliable across repeated trials (Figure 2B).

**Figure 2:**
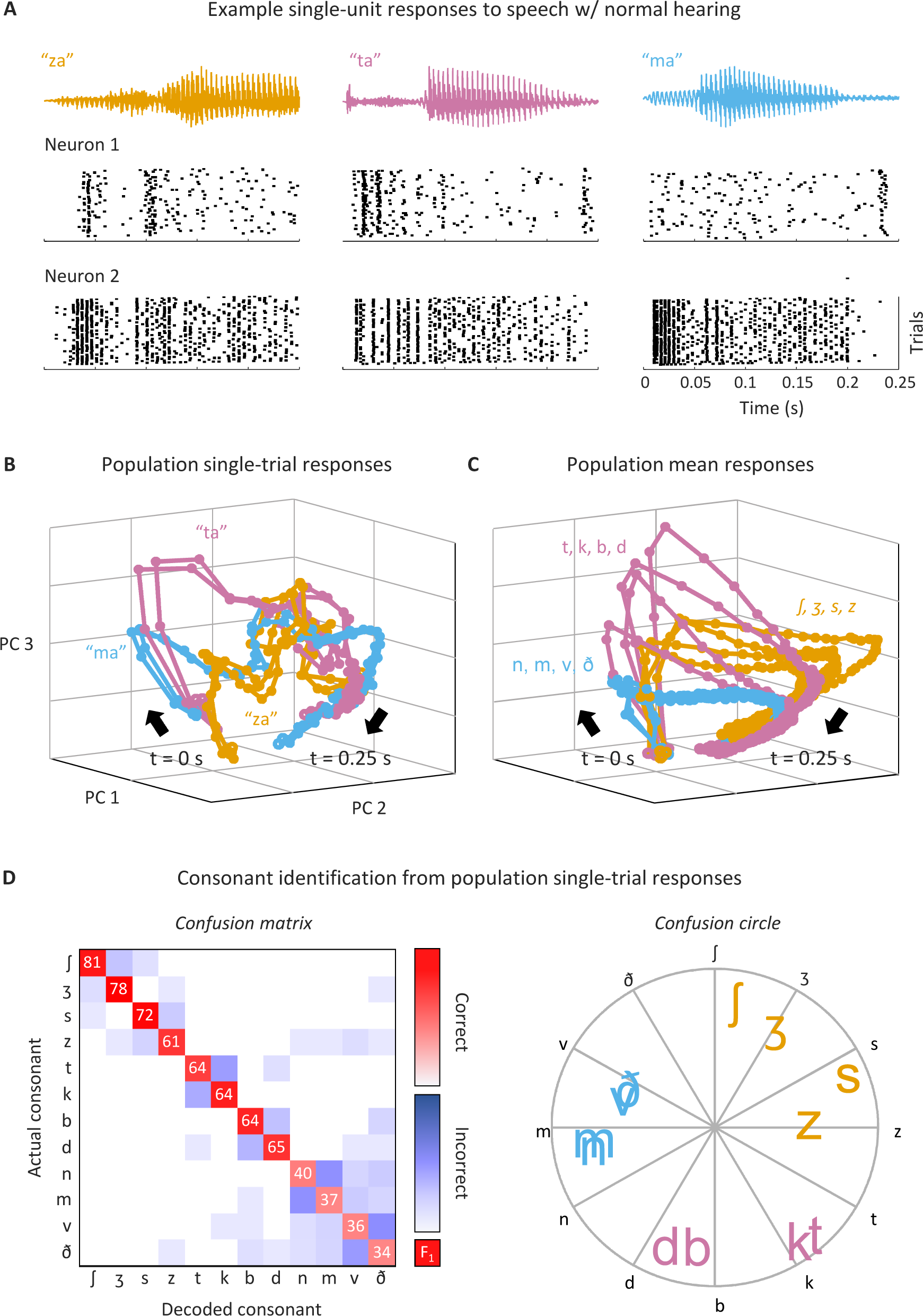
Single-trial responses to speech can be classified with high accuracy. (A) Single-unit responses to speech. Each column shows the sound waveform for one instance of a syllable and the corresponding raster plots for repeated presentations of that syllable for two example neurons from an animal with normal hearing. (B) Low-dimensional visualization of population single-trial responses to speech. Each line shows the responses from all neurons from all animals with normal hearing after principal component decomposition and projection into the space defined by the first three principal components. Responses to two repeated presentations for each of three syllables (indicated by the three colors) are shown. The time points corresponding to syllable onset are indicated by t = 0 s. (C) Low-dimensional visualization of mean population response to each consonant. Each line shows responses as in (B) after averaging across all presentations of syllables with the same consonant. Mean responses to each of 12 consonants are shown, with colors corresponding to consonant categories: sibilant fricatives (orange), stops (pink), and nasals and non-sibilant fricatives (blue). (D) Performance and confusion patterns for classification of single-trial responses. Left: Each row shows the frequency with which responses to one consonant were identified as that consonant (diagonal entries) or other consonants (off-diagonal entries) by the classifier. The values on the diagonal entries are the F1 score computed as 2 x (precision x recall) / (precision + recall), where precision = true positives / (true positives + false positives) and recall = true positives / (true positives + false negatives). The values shown are the average across all populations. For all consonant identification analyses shown in the Results, a support vector machine was used to classify the first 150 ms of single-trial responses of populations of 150 neurons represented as spike counts with 5 ms time bins. Right: Consonants were assigned angles along a unit circle (indicated by black letters). For each single-trial response for a given actual consonant, a vector was formed with magnitude 1 and angle corresponding to the consonant that the response was identified as by the classifier. The positions of the colored letters indicate the sum of these vectors across all responses for each consonant.

To assess the degree to which the neural code allowed for accurate identification of consonants, we trained a support vector machine classifier to identify the consonant in each syllable based on the population response patterns. Despite the variability in the responses to each consonant across syllables with different vowels and talkers, the average responses to different consonants were still distinct (Figure 2C) and single trials could be classified with high accuracy (Figure 2D). The errors made by the classifier reflected confusions within consonant classes as expected from human perceptual studies (Miller and Nicely, 1955; Phatak and Allen, 2007): accuracy was high for the sibilant fricatives (*∫, 3, s, z*), moderate for the stops (*t, k, b, d*), and low for the nasals (*n, m*) and the non-sibilant fricatives (*v, ð*).

We presented the same set of syllables to animals with hearing loss before and after processing with the hearing aid. The mean spike rate of individual neurons was decreased by hearing loss but restored to normal by the hearing aid (Figures 3A,B; for full details of all statistical tests including sample sizes and p-values, see Table S1). A classifier trained to detect speech in silence based on the neural response patterns of individual neurons confirmed that the hearing aid restored audibility to normal (Figure 3C). Consonant identification was also impacted by hearing loss but, unlike audibility, remained well below normal even with the hearing aid (Figure 3D). The hearing aid failed to restore consonant identification not only for speech in quiet, but also for speech presented in the presence of either a second independent talker or multi-talker noise. This failure was evident across a range of different classifiers, neural representations, and population sizes (Figures S1,S2) and, thus, reflects a general deficit in the neural code.

**Figure 3:**
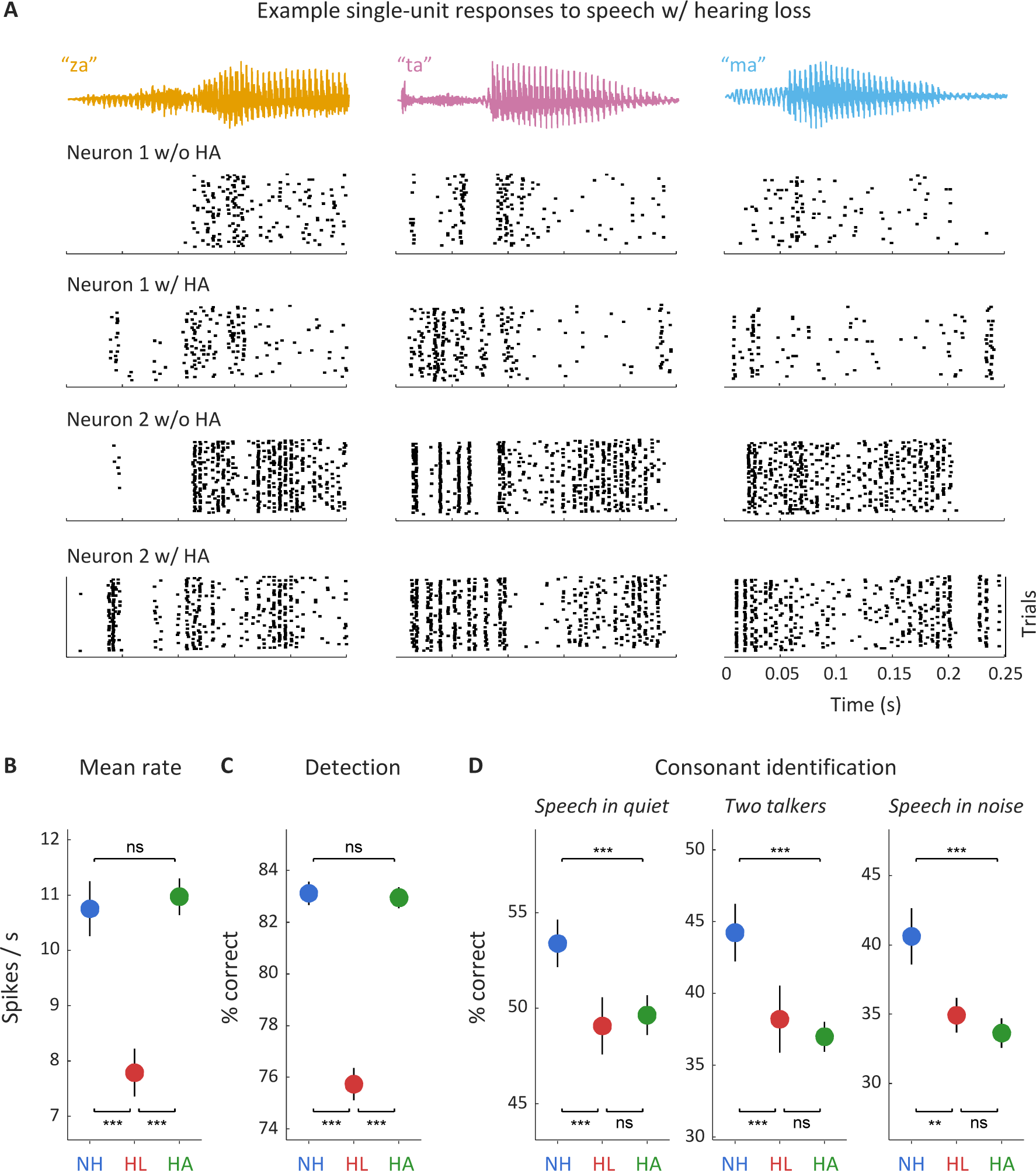
Hearing aids restore speech audibility but not consonant identification. (A) Single-unit responses to speech. Each column shows the sound waveform for one instance of a syllable and the corresponding raster plots for repeated presentations of that syllable for two example neurons from an animal with hearing loss, without and with a hearing aid. (B) Spike rate of single-unit responses to speech at 62 dB SPL. Results are shown for neurons from normal hearing animals (NH) and animals with hearing loss without (HL) and with (HA) a hearing aid (mean ± 95% confidence intervals derived from bootstrap resampling; for sample sizes and details of statistical tests for all figures, see Table S1). (C) Performance of a classifier trained to detect speech at 62 dB SPL in silence based on individual single-unit responses. (D) Performance of a classifier trained to identify consonants based on population responses to speech at 62 dB SPL. Results are shown for three conditions: speech in quiet, speech in the presence of ongoing speech from a second talker at equal intensity, and speech in the presence of multi-talker babble noise at equal intensity.

### Hearing aids fail to restore the selectivity of responses to speech

To understand why the hearing aid failed to restore consonant identification to normal, we investigated how different features of the neural response patterns varied across hearing conditions. Accurate auditory perception requires the response patterns elicited by different sounds to be distinct and reliable. For consonant identification, the response to a particular instance of a consonant must be similar to responses to other instances of that consonant but different from responses to other consonants.

In the context of any perceptual task, a neural response pattern can be separated into signal and noise, i.e. the components of the response which are helpful for the task and the components of the response which are not (Figure 4A). For consonant identification, the signal can be further divided into a *common signal*, which is common to all consonants, and a *differential signal*, which is specific to each consonant. The common signal reflects the average detectability (i.e. audibility) of all consonants, while the differential signal determines how well different consonants can be discriminated.

**Figure 4:**
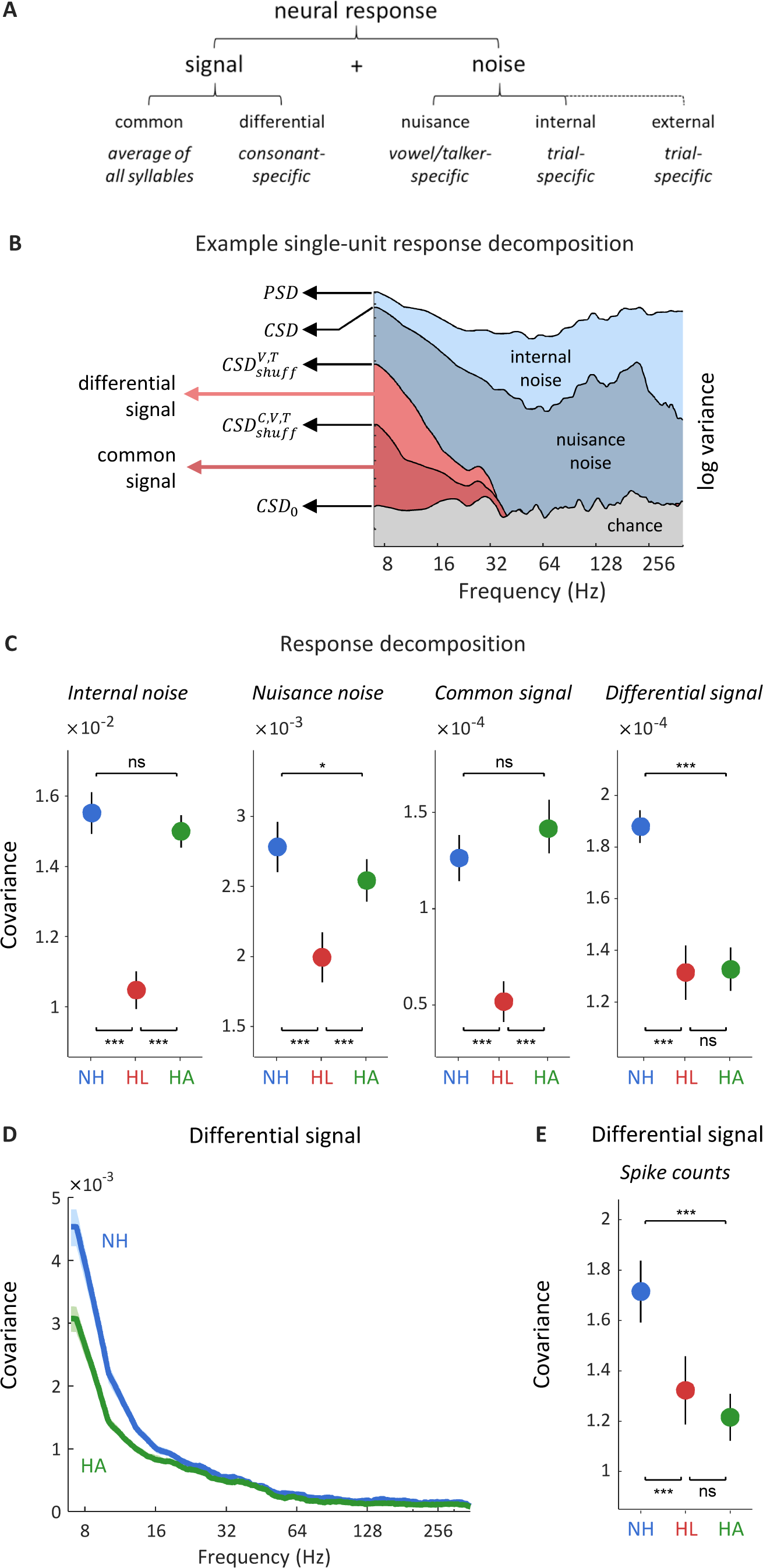
Hearing aids fail to restore the selectivity of neural responses to speech. (A) The different signal and noise components of neural responses in the context of a consonant identification task. (B) Spectral decomposition of responses for an example neuron. Each line shows a spectral density computed from responses before and after different forms of shuffling, and each filled area indicates the fraction of the total response variance corresponding to each response component. (C) Magnitude of different response components for single-unit responses to speech at 62 dB SPL. Results are shown for neurons from normal hearing animals (NH) and animals with hearing loss without (HL) and with (HA) a hearing aid (mean ± 95% confidence intervals derived from bootstrap resampling). (D) Magnitude of the differential signal component as a function of frequency for single-unit responses to speech at 62 dB SPL. Results are shown for neurons from normal hearing animals (NH) and animals with hearing loss with (HA) a hearing aid (mean ± 95% confidence intervals derived from bootstrap resampling indicated by shaded regions). (E) Magnitude of the differential signal component for single-unit spike counts in response to speech at 62 dB SPL.

The noise can also be further divided based on the different sources of variability in neural response patterns. The first source of variability is *nuisance noise*, which arises because consonants are followed by different vowels or spoken by different talkers (note that while this component of the response serves as noise for this task, it could also serve as signal for a different task, e.g. talker identification). The second source of variability is *internal noise*, which reflects the fundamental limitations on neural coding due to the stochastic nature of spiking and other intrinsic factors. For speech in the presence of additional sounds such as background noise, there is also *external noise*, which is the variability in responses that is caused by the additional sounds themselves.

All of these signal and noise components have the potential to influence consonant identification through their impact on the neural response patterns and together they form a complete description of any response. To isolate each of these components in turn, we computed the covariance between response patterns with different forms of shuffling across consonants, vowels, and talkers. We performed this decomposition of the responses in the frequency domain by computing spectral densities in order to gain further insight into which features of speech were reflected in each component.

The results are shown for a typical neuron for speech in quiet in Figure 4B. We first isolated the internal noise by comparing the power spectral density (*PSD*) of responses across a single trial of every syllable with the cross spectral density (*CSD*) of responses to repeated trials of the same speech (i.e. with the order of consonants, vowels, and talkers preserved). The *PSD* provides a frequency-resolved measure of the variance in a single neural response, while the *CSD* provides a frequency-resolved measure of the covariance between two responses. For an ideal neuron, repeated trials of identical speech would elicit identical responses and the *CSD* would be equal to the *PSD*. For a real neuron, the difference between the *PSD* and the *CSD* gives a measure of the internal noise. For the example neuron, the *CSD* was less than the *PSD* at all frequencies, but while the *PSD* was relatively flat the *CSD* decreased with increasing frequency up to 80 Hz and then increased between 80 and 250 Hz, indicating that the internal noise was smallest (and, thus, the neural responses most reliable) at frequencies corresponding to the envelope and voice pitch of the speech.

We next isolated the nuisance noise by comparing the *CSD* to the cross spectral density of responses to repeated trials after shuffling across vowels and talkers (denoted as 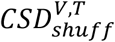). After this shuffling, the only remaining covariance between the responses is that which is shared across different instances of the same consonants. For the example neuron, this covariance was only significant at frequencies corresponding to the speech envelope; at frequencies higher than 40 Hz, the 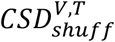 dropped below chance (denoted as *CSD*_0_). Thus, the nuisance noise, given by the difference between the *CSD* and the 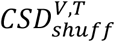, was largest at the frequencies corresponding to pitch (which is expected because pitch is reliably encoded in the responses but is not useful for talker-independent consonant identification).

Finally, we isolated the common signal from the differential signal by comparing the 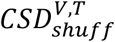 with the cross spectral density of the responses after shuffling across talkers, vowels, and consonants (denoted as 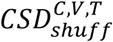). The only covariance between the responses that remains after this shuffling is that which is shared across all syllables. For the example neuron, both the differential signal, given by the difference between the 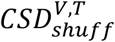 and the 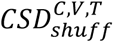, and the common signal, given directly by the 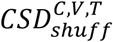, were significant across the full range of speech envelope frequencies.

At the population level, hearing loss impacted all components of the responses, with internal noise, nuisance noise, common signal, and differential signal all decreasing in magnitude (Figure 4C). The hearing aid increased the magnitude of the noise components but both the internal noise and the nuisance noise remained at or below normal levels. This suggests that mild-to-moderate hearing loss does not result in either fundamental limitations on neural coding or increased sensitivity to uninformative features of speech that can account for the failure of the hearing aid to restore consonant identification to normal.

The hearing aid also restored the common signal to normal, but failed to increase the magnitude of the differential signal. This suggests that the key difference between normal and aided responses is their selectivity, i.e. the degree to which their average responses to different consonants are distinct. This difference was most pronounced in the low-frequency component of the responses (Figure 4D); in fact, the same failure of the hearing aid to increase the differential signal was evident when looking only at spike counts (Figure 4E).

### The selectivity of aided responses to tones is normal

One possible explanation for the low selectivity of aided responses to speech is broadened frequency tuning, which would decrease sensitivity to differences in the spectral content of different consonants and increase the degree to which features of speech at one frequency are susceptible to masking by noise at other frequencies. The width of cochlear frequency tuning can increase with cochlear damage (Liberman et al., 1986) and impaired frequency selectivity is often reported in people with hearing loss (Moore, 2007). However, the degree to which frequency tuning is broadened with hearing loss depends on both the severity of the hearing loss and the intensity of incoming sounds (because frequency tuning broadens with increasing intensity even with normal hearing). Forward-masking paradigms that provide psychophysical estimates that closely match neural tuning curves (Moore and Glasberg, 1981; Shera et al., 2002; Sumner et al., 2018) suggest changes in frequency tuning may not be significant for mild-to-moderate hearing loss at moderate sound intensities (Nelson, 1991).

To characterize frequency tuning, we examined responses to pure tones presented at different frequencies and intensities. We defined the characteristic frequency (CF) of each neuron as the frequency that elicited a significant response at the lowest intensity and the threshold as the minimum intensity required to elicit a significant response at the CF (Figure 5A). Hearing loss caused an increase in thresholds across the range of speech-relevant frequencies, but this threshold shift was corrected by the hearing aid; in fact, aided thresholds were lower than those for normal hearing for CFs at both edges of the speech-relevant range (Figure 5B).

**Figure 5:**
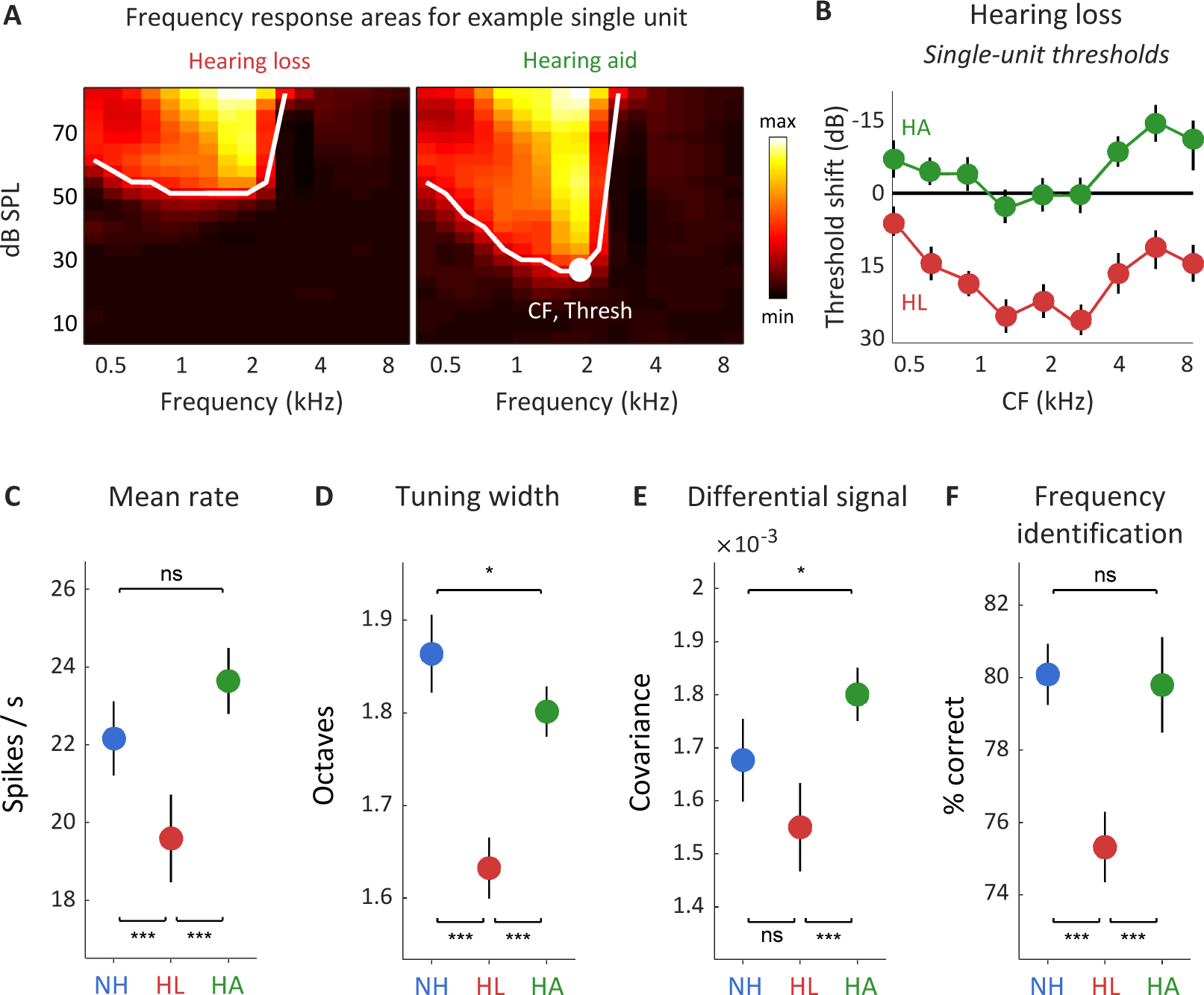
Hearing aids restore the selectivity of neural responses to tones. (A) Single-unit responses to tones. The FRA for an example single-unit from an animal with hearing loss showing the mean spike rate during the presentation of tones with different frequencies and intensities without (left) and with (right) a hearing aid. The center frequency and threshold are indicated. The lines indicate the lowest intensity for each frequency at which the response was significantly greater than responses recorded during silence (p < 0.01 for Poisson-distributed spike counts). (B) Threshold shift as a function of frequency for single-unit responses to tones. Results are shown for neurons from animals with hearing loss without (HL) and with (HA) a hearing aid. The values shown are the threshold shift relative to the mean of all neurons from all animals with normal hearing (mean ± 95% confidence intervals derived from bootstrap resampling). (C) Spike rate of single-unit responses to tones at 62 dB SPL. Results are shown for neurons from normal hearing animals (NH) and animals with hearing loss without (HL) and with (HA) a hearing aid. (D) Tuning width of single-unit responses to tones at 62 dB SPL. The values shown are the range of frequencies for which the mean spike rate during the presentation of a tone was at least half of its maximum value across all frequencies. (E) Magnitude of the differential signal component for single-unit responses to tones at 62 dB SPL. (F) Performance of a classifier trained to identify tone frequency based on population responses to tones at 62 dB SPL. A support vector machine was used to classify the first 150 ms of single-trial responses of populations of 10 neurons represented as spike counts with 5ms time bins.

The mean spike rate of individual neurons in response to pure tones presented at the same intensity as the speech (62 dB SPL) was decreased by hearing loss, but restored to normal by the hearing aid (Figure 5C). The width of frequency tuning at this intensity (defined as the range of frequencies for which the mean spike rate was at least half of its maximum value) was also decreased by hearing loss (Figure 5D), as expected given the increased thresholds. Tuning width was increased by the hearing aid, but remained slightly narrower than normal. This result suggests that mild-to-moderate hearing loss does not result in broadened frequency tuning at moderate intensities even after amplification by the hearing aid.

To determine directly whether the selectivity of responses to pure tones was impacted by hearing loss, we again isolated the differential signal component of the response. The magnitude of the differential signal was unimpacted by hearing loss and was slightly higher than normal with the hearing aid (Figure 5E), indicating that there was no loss of selectivity. To confirm the normal selectivity of aided responses to tones, we trained a classifier to identify tone frequencies based on neural response patterns. The performance of the classifier was decreased by hearing loss but returned to normal with the hearing aid (Figure 5F). Thus, the failure of the hearing aid to restore consonant identification to normal does not appear to result from a general loss of selectivity in neural responses.

### Hearing aid compression decreases the selectivity of responses to speech

Our results thus far suggest that if the low selectivity of aided responses to speech reflects a supra-threshold auditory processing deficit with hearing loss, it is only manifest for complex sounds. While this is certainly possible given the nonlinear nature of auditory processing, there is also another potential explanation: the low selectivity of responses to speech may be a result of distortions caused by the hearing aid itself (Kates, 2010; Souza, 2002). The multi-channel WDRC algorithm in the hearing aid constantly adjusts the amplification across frequencies, with each frequency receiving more amplification when it is weakly present in the incoming sound and less amplification when it is strongly present. This results in a compression of incoming sound across frequencies and time into a reduced range. Since a pure tone is a simple sound with a single frequency and constant amplitude, this compression has relatively little impact. But for complex sounds with multiple frequencies that vary in amplitude over time, such as speech, this compression serves to decrease both spectral and temporal contrast.

The WDRC algorithm is designed to replace the normal amplification and compression that are lost because of cochlear damage. But there are two potential problems with this approach. First, while normal cochlear compression does serve to decrease spectral and temporal contrast, there are also other mechanisms acting in a healthy cochlea that counteract this by increasing contrast -- e.g. cross-frequency suppression -- that are not included in the WDRC algorithm (Lesica, 2018). Second, there is evidence to suggest that with mild-to-moderate hearing loss, amplification of low intensity sounds is impaired but compression of moderate and high intensity sounds remains normal (Dubno et al., 2007; Lopez-Poveda et al., 2005; Plack et al., 2004). Thus, the total compression for the aided condition with mild-to-moderate hearing loss may be higher than normal, resulting in an effective decrease in the spectral and temporal contrast of complex sounds as represented in the neural code.

To investigate the impact of the hearing aid compression on the selectivity of responses to speech, we first computed the spectrograms of each instance of each consonant before and after processing with the hearing aid and measured their contrast (Figure 6A). On average, the spectrotemporal contrast after processing with the hearing aid was 15% lower than in the original sound (Figure 6B). This decrease in contrast was reflected in the performance of a classifier trained to identify the consonant in each spectrogram, which also decreased after processing with the hearing aid (Figure 6C).

**Figure 6:**
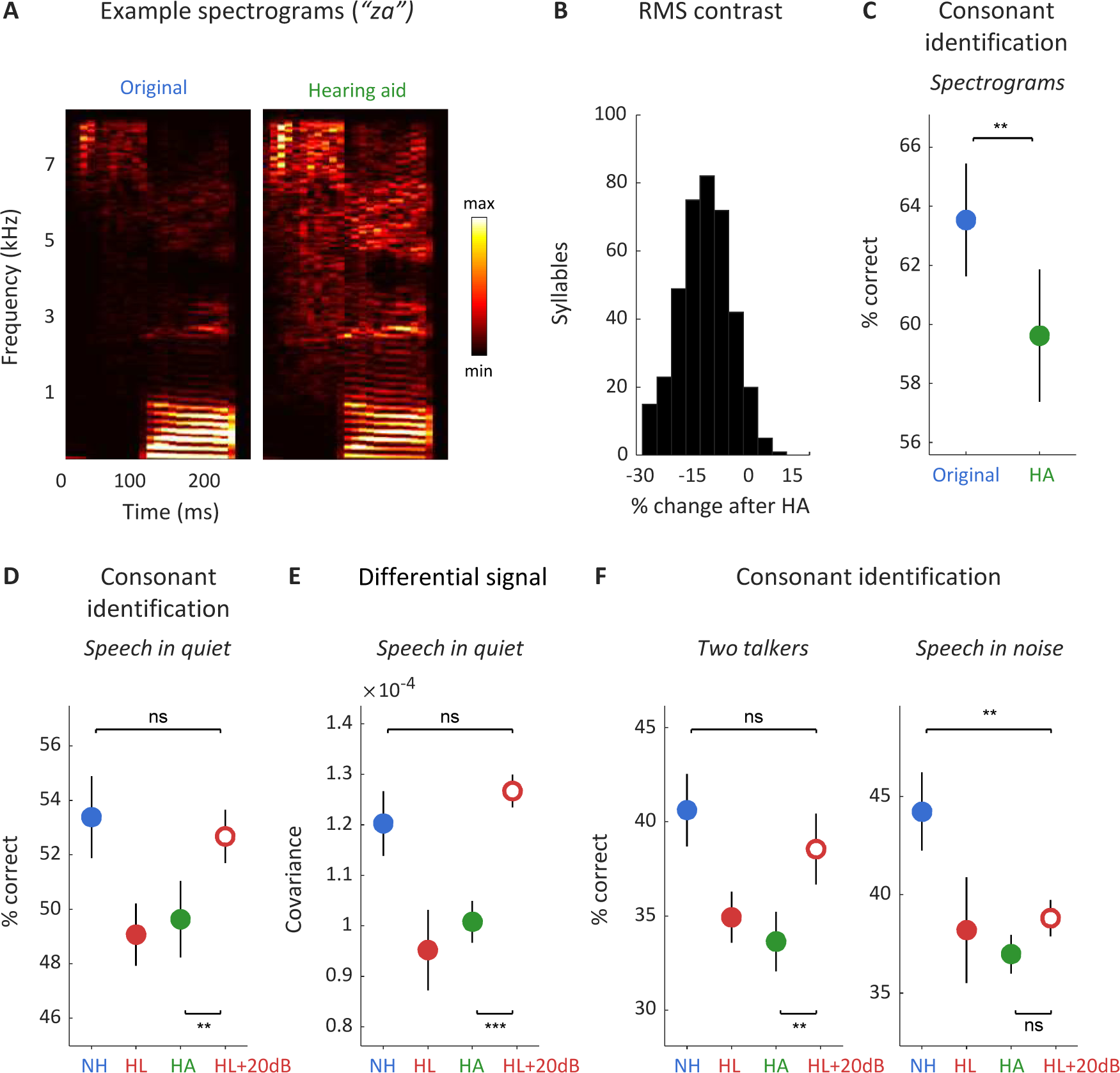
Hearing aid compression decreases the selectivity of neural responses to speech. (A) Spectrograms showing the log power across frequencies at each time point in one instance of the syllable “za” before and after processing with a hearing aid. (B) Percent change in RMS contrast of all syllables (n = 384) after processing with a hearing aid. Only the first 150 ms of each syllable were used. (C) Performance of a support vector machine classifier trained to identify consonants based on spectrograms (mean ± standard error across 10 different held-out samples). (D) Performance of a classifier trained to identify consonants based on population responses to speech at 62 dB SPL. Results are shown for normal hearing animals (NH) and animals with hearing loss without a hearing aid (HL), with a hearing aid (HA), and with linear amplification (HL+20dB) (mean ± 95% confidence intervals derived from bootstrap resampling). (E) Magnitude of the differential signal component for single-unit responses to speech at 62 dB SPL. (F) Performance of a classifier trained to identify consonants based on population responses to speech at 62 dB SPL. Results are shown for speech in the presence of ongoing speech from a second talker at equal intensity and speech in the presence of multi-talker babble noise at equal intensity.

If the hearing aid compression is responsible for the low selectivity of neural responses, then it should be possible to improve selectivity (and, thus, consonant identification) by providing amplification without compression. We presented the same consonant-vowel syllables after linear amplification (with a fixed gain of 20 dB) and compared the results of classification and response decomposition to those for the original speech. Linear amplification without compression restored both classifier performance and the magnitude of the differential signal to normal (Figures 6D,E). Thus, the failure of the hearing aid to restore response selectivity and consonant identification in quiet appears to result from hearing aid compression rather a deficit in supra-threshold auditory processing with hearing loss. Linear amplification is able to restore the selectivity of neural responses and, consequently, consonant identification by restoring audibility without distorting the spectral and temporal features of speech.

### Amplification decreases consonant identification in noise for all hearing conditions

We next investigated whether removing hearing aid compression and providing only linear amplification was also sufficient to restore consonant identification to normal in the presence of additional sounds. While linear amplification was sufficient to restore consonant identification in the presence of a second independent talker, it failed in multi-talker noise (Figure 6F). This suggests that for speech in noise, there are additional reasons for the failure of the hearing aid to restore consonant identification beyond just the distortions caused by hearing aid compression.

The failure of both the hearing aid and linear amplification to restore consonant identification in noise could reflect a supra-threshold auditory processing deficit with hearing loss that is only manifest in difficult listening conditions, but this is not necessarily the case. Even with normal hearing, the intelligibility of speech in noise decreases as overall intensity increases (an effect known as ‘rollover’ with a complex physiological basis (Dubno et al., 2012; Horvath and Lesica, 2011; Wong et al., 1998)). When the background noise is dominated by low frequencies (as is the case for multi-talker noise), speech intelligibility decreases by approximately 5% for every 10 dB increase in overall intensity above moderate levels, even when the speech-to-noise ratio remains constant (Hornsby et al., 2005; Studebaker et al., 1999). Thus, the differences in the perception of moderate-intensity speech-in-noise with normal hearing and that of amplified speech-in-noise with hearing loss may not reflect the effects of hearing loss per se, but rather the unintended consequences of amplifying sounds to high intensities to restore audibility.

To assess the impact of rollover on the neural code, we compared consonant identification and response decomposition with normal hearing before and after linear amplification. The amplification to high intensity did not impact consonant identification in quiet or in the presence of second talker, but decreased consonant identification in multi-talker noise (Figure 7A). This decrease in consonant identification in noise at high intensities with normal hearing appears to result from a decrease in response selectivity; the magnitude of the differential signal was significantly smaller after amplification, while the magnitudes of the common signal and total noise were unchanged (Figure 7B; note that because we did not present repeated trials of ‘frozen’ multi-talker noise, we cannot isolate the individual noise components but we can still measure the total magnitude of all noise components as the difference between the *PSD* and the 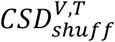).

**Figure 7:**
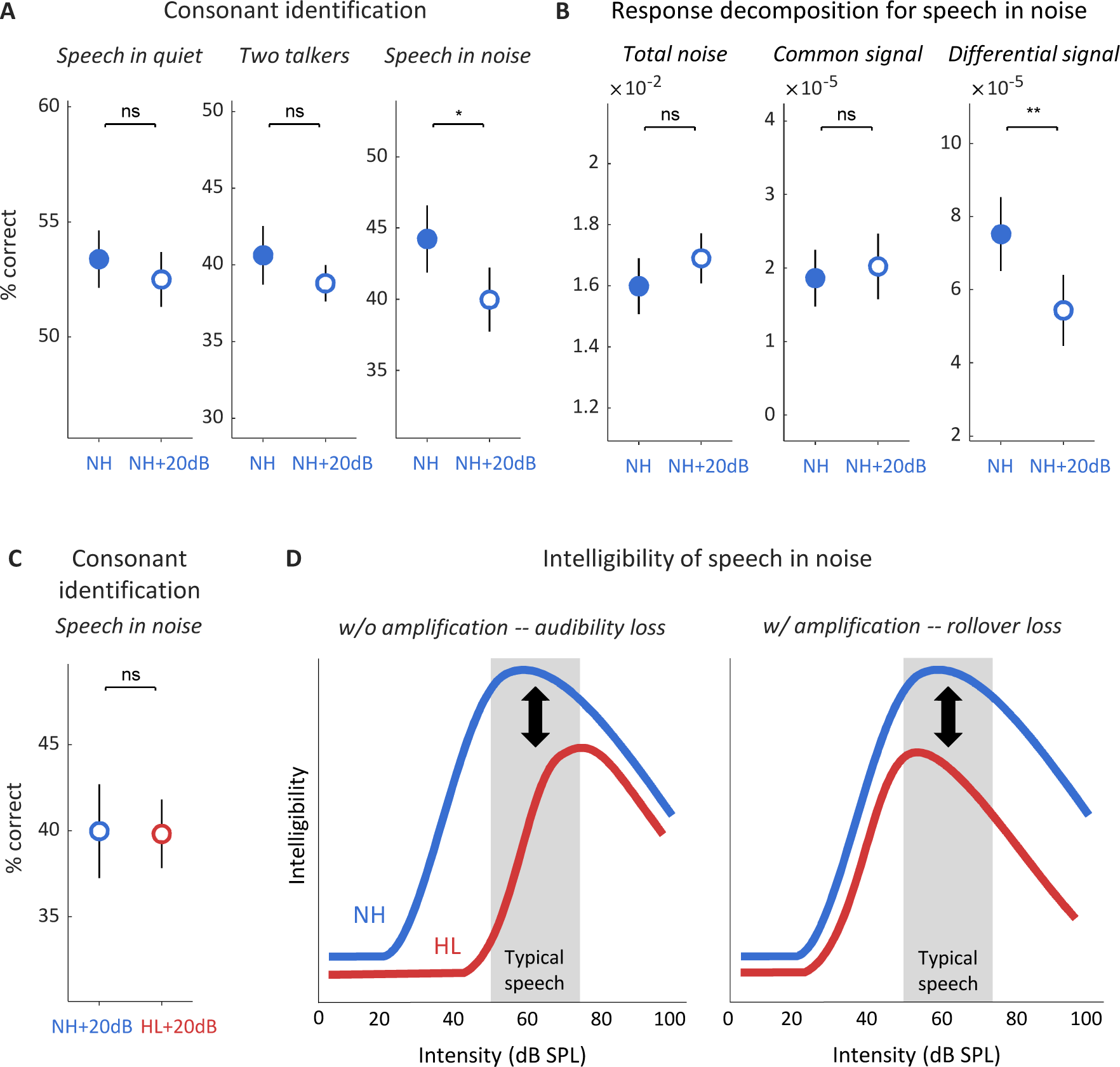
Amplification decreases consonant identification even with normal hearing. (A) Performance of a classifier trained to identify consonants based on population responses to speech at 62 dB SPL. Results are shown for normal hearing animals without (NH) and with (NH+20dB) linear amplification (mean ± 95% confidence intervals derived from bootstrap resampling) for three conditions: speech in quiet, speech in the presence of ongoing speech from a second talker at equal intensity, and speech in the presence of multi-talker babble noise at equal intensity. (B) Magnitude of different response components for single-unit responses to speech at 62 dB SPL in multi-talker babble noise. (C) Performance of a classifier trained to identify consonants based on population responses to speech at 62 dB SPL in noise with linear amplification. Results are shown for normal hearing animals (NH+20dB) and animals with hearing loss (HL+20dB). (D) A schematic diagram showing the effects of intensity on speech intelligibility with and without hearing loss and amplification. The range of intensities of typical speech is shown in gray. Left: The loss of intelligibility with hearing loss that results from the loss of audibility without amplification. Right: The loss of intelligibility with hearing loss that results from rollover with amplification.

To determine whether rollover can account for the deficit in consonant identification in noise with hearing loss that remains even after linear amplification, we compared consonant identification after linear amplification for both hearing loss and normal hearing (i.e. using responses to amplified speech for both conditions). When compared at the same high intensity, consonant identification with or without hearing loss was not significantly different (Figure 7C). Thus, the failure of both the hearing aid and linear amplification to restore consonant identification in noise does not appear to reflect a deficit in supra-threshold processing caused by hearing loss, but rather a deficit in high-intensity processing that is present even with normal hearing.

Taken together, our results provide a clear picture of the challenge that must be overcome to restore normal auditory perception after mild-to-moderate hearing loss. Amplification is required to restore audibility, but can also reduce the selectivity of neural responses in complex listening conditions. Thus, a hearing aid must provide amplification while also transforming incoming sounds to compensate for the loss of selectivity at high intensities. Current hearing aids provide the appropriate amplification but fail to implement the required additional transformation and, in fact, appear to further decrease selectivity through compression that decreases the spectrotemporal contrast of incoming sounds.

## Discussion

This study was designed to identify the reasons why hearing aids fail to restore normal auditory perception through analysis of the underlying neural code. Our results suggest that difficulties during aided listening with mild-to-moderate hearing loss arise primarily from the low selectivity of neural responses. While a hearing aid corrected many of the changes in neural response patterns that were caused by hearing loss, the average response patterns elicited by different consonants remained much less distinct than with normal hearing. The low selectivity of aided responses to speech did not appear to reflect a fundamental deficit in supra-threshold auditory processing as the selectivity of responses to moderate-intensity tones was normal. In fact, for speech in quiet, the low selectivity resulted from compression in the hearing aid itself that decreased the spectrotemporal contrast of incoming sounds; linear amplification without compression restored selectivity and consonant identification to normal. For speech in multi-talker noise, however, selectivity and consonant identification remained low even after linear amplification. But linear amplification also decreased the selectivity of neural responses with normal hearing such that, when compared at the same high intensity, consonant identification in noise with normal hearing and hearing loss were similar.

These results are consistent with the idea that for mild-to-moderate hearing loss, decreased speech intelligibility is primarily caused by decreased audibility (Zurek and Delhorne, 1987) rather than supra-threshold processing deficits. Of course, there are many listeners whose problems go beyond audibility: more severe or specific hearing loss may result in additional supra-threshold deficits (Moore, 2007); cognitive factors may interact with hearing loss to create additional problems (Humes and Dubno, 2010); and supra-threshold deficits can exist without any significant loss of audibility for a variety of reasons (Parthasarathy et al., 2020). But numerous perceptual studies have reported that the intelligibility of speech-in-noise at high intensities for people with mild-to-moderate hearing loss is essentially normal (Ching et al., 1998; Lee and Humes, 1993; Oxenham and Kreft, 2016; Studebaker et al., 1999; Summers and Cord, 2007). Unfortunately, because of rollover, even normal processing is impaired at high intensities. Thus, those with hearing loss must currently choose between listening naturally to low- and moderate-intensity sounds and suffering from reduced audibility, or artificially amplifying sounds to high intensities and suffering from rollover (Figure 7D).

Overcoming the current tradeoff between loss of audibility and rollover is a challenge, but our results are encouraging with respect to the potential of future hearing aids to bring significant improvements. We found that current hearing aids already restore many aspects of the neural code for speech to normal, including mean spike rates, selectivity for pure tones, fundamental limitations on coding (as reflected by internal noise), and sensitivity to prosodic aspects of speech (as reflected by nuisance noise). Instead of compression, which appears to exacerbate the loss of selectivity that accompanies amplification to high intensities, the next-generation of hearing aids must incorporate additional processing to counteract the mechanisms that cause rollover. There have been a number of previous attempts to manipulate the features of speech to improve perception by, for example, enhancing spectral contrast (Baer et al., 1993; May et al., 2018; Miller et al., 1999; Moore, 2016; Rallapalli and Alexander, 2019; Rasetshwane et al., 2014). But these strategies have typically been developed to counteract processing deficits that are a direct result of severe hearing loss, e.g. loss of cross-frequency suppression, that may not be present with mild-to-moderate loss. New approaches that are specifically designed to improve perception at high intensities even for normal hearing listeners may be more effective.

The multi-channel WDRC algorithm in current hearing aids is designed to compensate for the dysfunction of outer hair cells (OHCs) in the cochlea. The OHCs normally provide amplification and compression of incoming sounds, but with hearing loss their function is often impaired either through direct damage or through damage to supporting structures (Gates and Mills, 2005). The true degree of OHC dysfunction in any individual is difficult to determine, so the WDRC algorithm provides amplification and compression in proportion to the measured loss of audibility across different frequencies, which reflects loss of amplification. But while severe hearing loss may result in a loss of both amplification and compression, several studies have found that mild-to-moderate hearing loss appears to result in a loss of amplification only (Dubno et al., 2007; Lopez-Poveda et al., 2005; Plack et al., 2004). Thus, with mild-to-moderate loss, the use of a WDRC hearing aid can result in excess compression that distorts the acoustic features of speech (Kates, 2010; Souza, 2002). Our results demonstrate that these distortions result in the representation of different speech elements in the neural code being less distinct from each other.

A number of studies of speech perception in people with mild-to-moderate hearing loss have found that linear amplification without compression is often comparable or superior to WDRC hearing aids (Davies-Venn et al., 2009; Kates, 2010; Larson et al., 2000; Shanks et al., 2002). Our analysis of the neural code provides a physiological explanation for these findings and adds support to the growing movement to increase uptake of hearing aids through the development and provision of simple, inexpensive devices that can be obtained over-the-counter (Committee on Accessible and Affordable Hearing Health Care for Adults et al., 2016; Warren and Grassley, 2017). (Note that linear amplification alone is not sufficient for real-world devices; there must also be limits to prevent high-intensity sounds from being further amplified to uncomfortable or dangerous levels). Recent clinical evaluations of over-the-counter personal sound amplification products (PSAPs) have shown that they often provide similar benefit to premium hearing aids fit by professional audiologists (Brody et al., 2018; Cho et al., 2019; Humes et al., 2017). Thus, there is now sufficient physiological, psychophysical, and clinical evidence to conclude that simple devices can provide benefit for people with mild-to-moderate hearing loss that is comparable to that provided by current state-of-the-art devices.

This conclusion has important implications for strategies to combat the global burden of hearing loss. Simple devices may only be appropriate for people with mild-to-moderate loss, but this group currently includes more than 500 million people worldwide (Wilson et al., 2017). Thus, the wide adoption of simple devices could have a substantial impact, especially in low- and middle-income countries where the burden of hearing loss is largest and the uptake of hearing aids is lowest. Ideally, the next generation of state-of-the-art hearing aids will bring improvements in both benefit and affordability. But given the need for urgent action to mitigate the impact of hearing loss on wellbeing and mental health (Livingston et al., 2017; Wilson et al., 2017) and the potential for simple devices to provide significant benefit, promoting their use should be considered as a potential public health priority.

## Materials and Methods

### Experimental protocol

Experiments were performed on 35 young-adult gerbils of both sexes that were born and raised in standard laboratory conditions. Twenty of the animals were exposed to noise when they were 10-12 weeks old. ABR recordings and large-scale IC recordings were made from all animals when they were 14-18 weeks old.

#### Noise exposure

Sloping mild-to-moderate sensorineural hearing loss was induced by exposing anesthetized gerbils to high-pass filtered noise with a 3 dB/octave roll-off below 2 kHz at 118 dB SPL for 3 hours (Suberman et al., 2011). For anesthesia, an initial injection of 0.2 ml per 100 g body weight was given with fentanyl (0.05 mg per ml), medetomidine (1 mg per ml), and midazolam (5 mg per ml) in a ratio of 4:1:10. A supplemental injection of approximately 1/3 of the initial dose was given after 90 minutes. Internal temperature was monitored and maintained at 38.7° C.

#### Auditory brainstem responses

Animals were placed in a sound-attenuated chamber, and anesthesia and internal temperature were maintained as for noise exposure. An ear plug was inserted into one ear and a free-field speaker was placed 10 cm from the other ear. The sound level was calibrated prior to each recording using a microphone that was placed next to the open ear. Subdermal needles were used as electrodes with the active electrode placed behind the open ear, the reference placed over the nose, and the ground placed in a rear leg. Recordings were bandpass filtered between 300 and 3000 Hz. Clicks (0.1 ms) and tones (4 ms with frequencies ranging from 500 Hz to 8000 Hz in1 octave steps with 0.5 ms cosine on and off ramps) were presented at intensities ranging from 5 dB SPL to 85 dB SPL in 5 dB steps with a 25 ms pause between presentations. All sounds were presented 2048 times (1024 times with each polarity). Thresholds were defined as the lowest intensity at which the RMS of the mean response across presentations was more than twice the RMS of the mean of 2048 trials of activity recorded during silence.

#### Large-scale electrophysiology

Animals were placed in a sound-attenuated chamber and anesthetized for surgery with an initial injection of 1 ml per 100 g body weight of ketamine (100 mg per ml), xylazine (20 mg per ml), and saline in a ratio of 5:1:19. The same solution was infused continuously during recording at a rate of approximately 2.2 µl per min. Internal temperature was monitored and maintained at 38.7° C. A small metal rod was mounted on the skull and used to secure the head of the animal in a stereotaxic device. Two craniotomies were made along with incisions in the dura mater, and a 256-channel multi-electrode array (Neuronexus) was inserted into the central nucleus of the IC in each hemisphere (Figure 1A). The arrays were custom-designed to maximize coverage of the portion of the gerbil IC that is sensitive to the frequencies that are present in speech.

#### Multi-unit activity

MUA was measured from recordings on each channel of the array as follows: (1) a high pass filter was applied with a cutoff frequency of 500 Hz; (2) the absolute value was taken; (3) a low pass filter was applied with a cutoff frequency of 300 Hz. This measure of multi-unit activity does not require choosing a threshold; it simply assumes that the temporal fluctuations in the power at frequencies above 500 Hz reflect the spiking of neurons near each recording site.

#### Spike sorting

Single-unit spikes were isolated using Kilosort (Pachitariu et al., 2016) with default parameters. Recordings were separated into overlapping 1-hour segments with a new segment starting every 15 minutes. Kilosort was run separately on each segment and clusters from separate segments were chained together if at least 90% of their events were identical during their period of overlap. Clusters were retained for analysis only if they were present for at least 2.5 hours of continuous recording. This persistence criterion alone was sufficient to identify clusters that also satisfied the usual single-unit criteria with clear isolation from other clusters, lack of refractory period violations, and symmetric amplitude distributions (see Figure S3).

#### Sounds

Sounds were delivered to speakers (Etymotic ER-2) coupled to tubes inserted into both ear canals along with microphones (Etymotic ER-10B+) for calibration. The frequency response of these speakers measured at the entrance of the ear canal was flat (± 5 dB SPL) between 0.2 and 8 kHz. The full set of sounds presented is described below. All sounds other than multi-talker speech babble noise were presented diotically.

1. *Tone set 1*: 50 ms tones with frequencies ranging from 500 Hz to 8000 Hz in 0.5 octave steps and intensities ranging from 6 dB SPL to 83 dB SPL in 7 dB steps with 2 ms cosine on and off ramps and 175 ms pause between tones. Tones were presented 8 times each in random order.
2. *Tone set 2*: 50 ms tones with frequencies ranging from 500 Hz to 8000 Hz in 0.5 octave steps at 62 dB SPL with 5 ms cosine on and off ramps and 175 ms pause between tones. Tones were presented 128 times each in random order.
3. *Consonant-vowel (CV) syllables*: Speech utterances taken from the Articulation Index LSCP (LDC Catalog No.: LDC2015S12). Utterances were from 8 American English speakers (4 male, 4 female). Each speaker pronounced CV syllables made from all possible combinations of 12 consonants and 4 vowels. The consonants included the sibilant fricatives *∫, 3, s*, and *z*, the stops *t, k, b*, and *d*, the nasals *n* and *m*, and the non-sibilant fricatives *v* and *ð*. The vowels included *a, æ, i*, and *o*. Utterances were presented in random order with 175 ms pause between sounds at an intensity of 62 dB SPL (or 82 dB SPL after 20 dB linear amplification).
4. *Second independent talker*: Speech from 16 different talkers taken from the UCL Scribe database (https://www.phon.ucl.ac.uk/resource/scribe) provided by Prof. Mark Huckvale was concatenated to create a continuous stream of ongoing speech with one talker at a time.
5. *Omni-directional multi-talker speech babble noise*: Speech from 16 different talkers from the Scribe database was summed to create speech babble. The speech from each talker was first passed through a gerbil head-related transfer function (Maki and Furukawa, 2005) using software provided by Dr. Rainer Beutelmann (Carl von Ossietzky University) to simulate its presentation from a random azimuthal angle.

#### Hearing aid simulation

10-channel WDRC processing was simulated using a program provided by Prof. Johsua Alexander (Purdue University) (Alexander and Masterson, 2015). The crossover frequencies between channels were 200, 500, 1000, 1750, 2750, 4000, 5500, 7000, and 8500 Hz. The intensity thresholds below which amplification was linear for each channel were 45, 43, 40, 38, 35, 33, 28, 30, 36, and 44 dB SPL. The attack and release times for all channels were 5 and 40 ms, respectively. The gain and compression ratio for each channel were fit individually for each ear of each animal using the Cam2B.v2 software provided by Prof. Brian Moore (Cambridge University) (Moore et al., 2010). The gain before compression typically ranged from 10 dB at low frequencies to 25 dB at high frequencies. The compression ratios typically ranged from 1 to 2.5.

### Data analysis

#### Visualization of population response patterns

To reduce the dimensionality of population response patterns, the responses for each neuron were first converted to spike count vectors with 5 ms time bins. The responses to all syllables from all neurons across all animals for a given hearing condition were combined into one matrix and a principal component decomposition was performed to find a small number of linear combinations of neurons that best described the full population. To visualize responses in three dimensions, single trial or mean responses were projected into the space defined by the first three principal components.

#### Classification of population response patterns

Populations were formed by taking random, non-overlapping samples of neurons from across all animals for a given hearing condition. (Note that each population thus contained both simultaneously and non-simultaneously recorded neurons. The simultaneity of recordings could impact classification if the responses contain noise correlations, i.e. correlations in trial-to-trial variability, which would be present only in simultaneous recordings. But we have shown previously under the same experimental conditions that the noise correlations in IC populations are negligible (Garcia-Lazaro et al., 2013). This was also true of the populations used in this study (Figure S4)).

For the analyses shown in the Results, populations of 150 neurons were used and classification was performed after converting the responses for each neuron to spike count vectors with 5 ms time bins. Only the first 150 ms of the responses to each syllable were used to minimize the influence of the vowel. The classifier was a support vector machine with a max-wins voting strategy based on all possible combinations of binary classifiers and 10-fold cross validation. To ensure the generality of the results, different classifiers, neural representations, and population sizes were also tested (see Figures S2,S3).

#### Computation of spectral densities

Spectral densities were computed as a measure of the frequency-specific covariance between two responses (or variance of a single response). To compute spectral densities, responses to all syllables with different consonants and vowels spoken by different talkers were concatenated in time and converted to binary spike count vectors with 1 ms time bins

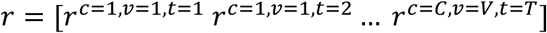

where *r*^*c,v,t*^ = [*r*^*c,v,t*^[1] *r*^*c,v,t*^[2] … *r*^*c,v,t*^[*N*]] is the binary spike count vector with *N* time bins for the response to one syllable composed of consonant *c* and vowel *v* spoken by talker *t*. Responses were then separated into 300 ms segments with 50% overlap and each segment was multiplied by a Hanning window. The cross spectral density between two responses was then computed as the average across segments of the discrete Fourier transform of one response with the complex conjugate of the discrete Fourier transform of the other response

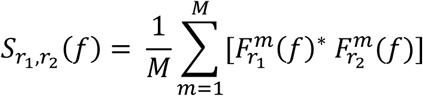

where 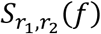 is the cross spectral density between responses *r*_1_ and *r*_2_, *M* is the total number of segments, 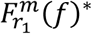 is the complex conjugate of the discrete Fourier transform of the *m*^*th*^ segment of *r*_1_, and 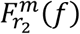 is the discrete Fourier transform of the *m*^*th*^ segment of *r*_2_. The values for negative frequencies were discarded. The final spectral density was smoothed using a median filter with a width of 0.2 octaves and scaled such that its sum across all frequencies was equal to the total covariance between the two responses

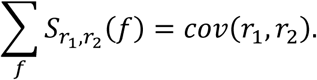

Several different spectral densities were computed before and after shuffling the order of the syllables in the concatenated responses to isolate different sources of covariance as described in the Results.

*PSD* - the power spectral density of a single response:

*r*_*1*_ = response to one trial of speech with all syllables in original order

*r*_2_ = *r*_1_

*CSD* - the cross spectral density of responses to repeated identical trials:

*r*_1_= response to one trial of speech with all syllables in original order

*r*_2_ = response to another trial of speech with all syllables in original order

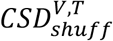 - the cross spectral density of responses after shuffling of vowels and talkers, leaving the responses matched for consonants only:

*r*_1_= response to one trial of speech with all syllables in original order

*r*_2_ = response to another trial of speech after shuffling of vowels and talkers

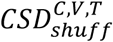 - the cross spectral density of responses after shuffling of consonants, vowels and talkers, leaving the responses matched for syllable onset only:

*r*_1_= response to one trial of speech with all syllables in original order

*r*_2_ = response to another trial of speech after shuffling of consonants, vowels and talkers

*CSD*_0_ - the cross spectral density of responses after shuffling of consonants, vowels and talkers and randomization of the phase of the Fourier transform of each response segment, leaving the responses matched for overall magnitude spectrum only:

*r*_1_= response to one trial of speech with all syllables in original order

*r*_2_ = response to another trial of speech after shuffling of consonants, vowels and talkers

To isolate the differential signal component of responses to tones, the same approach was used with shuffling of frequencies.

#### Classification of spectrograms

To convert sound waveforms to spectrograms, they were first separated into 80 ms segments with 87.5% overlap, then multiplied by a Hamming window. The discrete Fourier transform of each segment was taken, then the magnitude was extracted and converted to a logarithmic scale. Classification was performed using a support vector machine as described above for neural responses. Only the first 150 ms of the responses to each syllable were used.

## Acknowledgements

This work was supported by a Wellcome Trust Senior Research Fellowship [200942/Z/16/Z]. The authors thank J. Linden, S. Rosen, D. Fitzpatrick, B. Moore, J. Alexander, M. Huckvale, K. Harris, G. Huang, T. Keck and R. Beutelmann for their advice.

---

**Figure S1:**
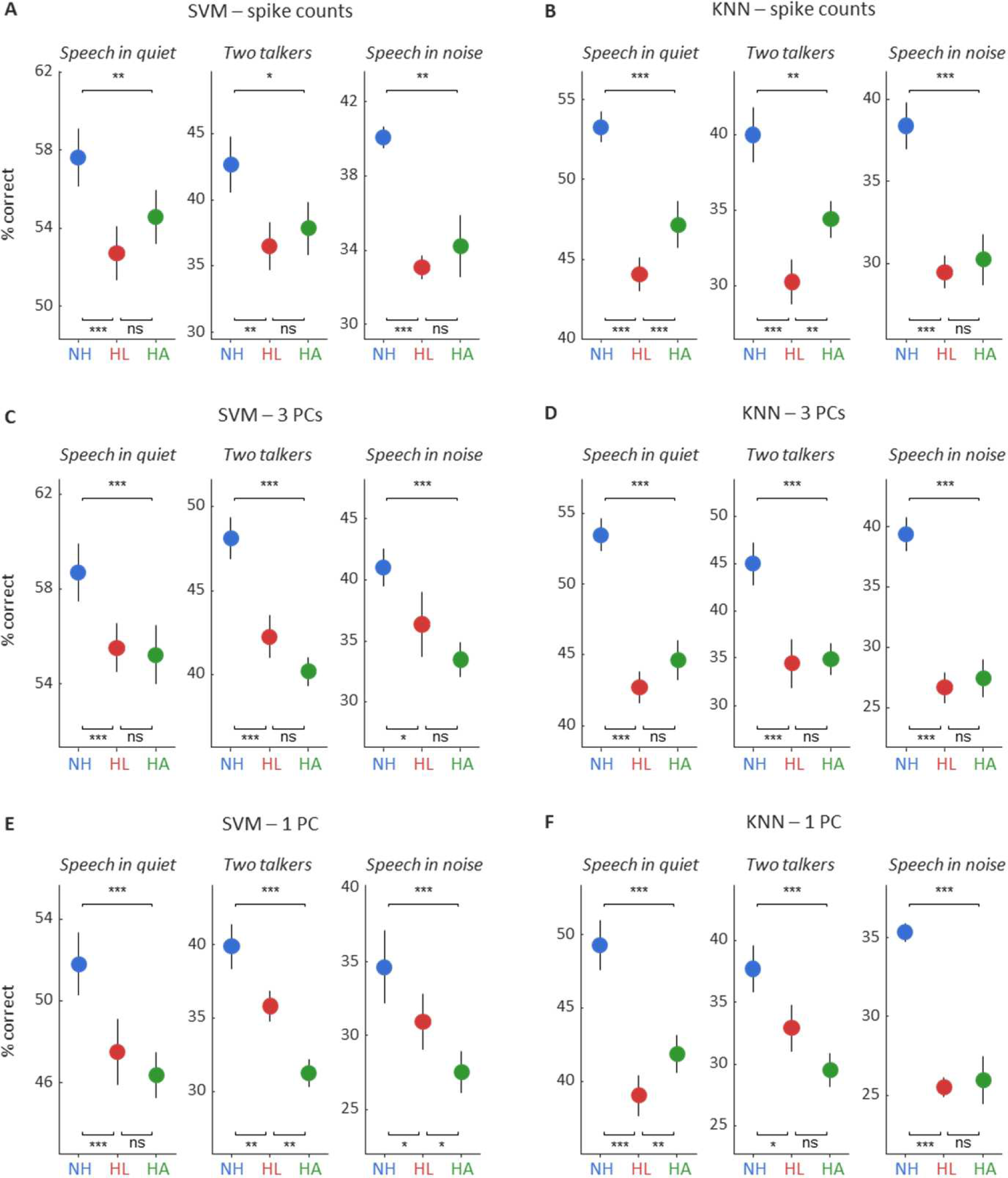
Consonant identification with different classifiers and neural representations. The figure shows the performance of different classifiers trained to identify consonants based on population responses to speech at 62 dB SPL with different response representations. In all cases, the first 150 ms of single-trial responses of populations of 150 neurons were used. For each classifier and neural representation, results are shown for three conditions: speech in quiet, speech in the presence of ongoing speech from a second talker at equal intensity, and speech in the presence of multi-talker babble noise at equal intensity. (A) Performance of a support vector machine trained to classify the total spike counts. The details of the support vector machine were identical to those used in the Results. (B) Performance of a k-nearest neighbors classifier trained to classify the total spike counts with 10-fold cross validation. The values shown are for k = 16 which had the highest cross-validated performance. (C) Performance of a support vector machine trained to classify the responses after projection onto the three principal components that best described the variance in responses across the entire population (reducing the response of the entire population to three values in each 5 ms time bin as in Figure 2B). (D) Performance of a k-nearest neighbors classifier trained to classify the responses after projection onto the first three principal components. (E) Performance of a support vector machine trained to classify the responses after projection onto the principal component that best described the variance in responses across the entire population (reducing the response of the entire population to one value in each 5 ms time bin). (F) Performance of a k-nearest neighbors classifier trained to classify the responses after projection onto the first principal component.

**Figure S2:**
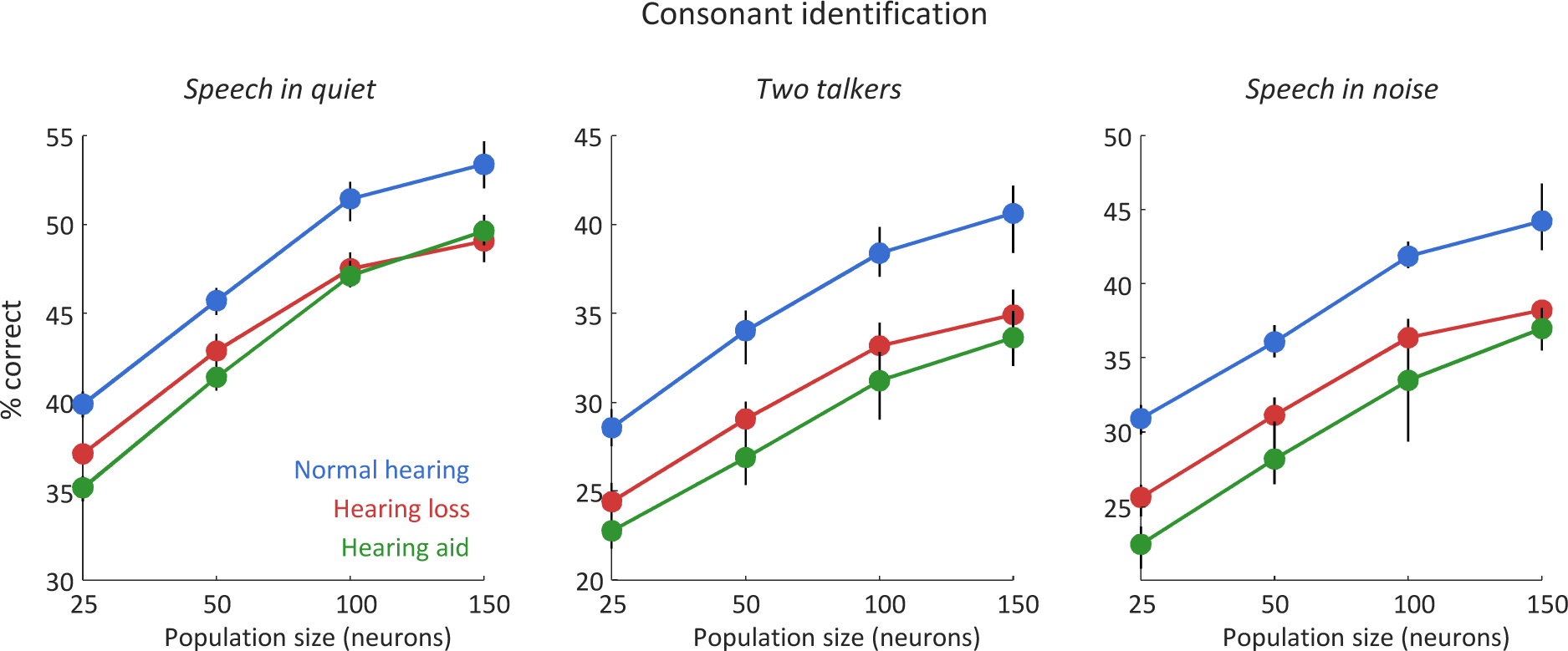
Consonant identification with different population sizes. The figure shows the performance of a support vector machine trained to identify consonants based on population responses to speech at 62 dB SPL as in the Results with different populations sizes (mean ± 95% confidence intervals derived from bootstrap resampling). Results are shown for three conditions: speech in quiet, speech in the presence of ongoing speech from a second talker at equal intensity, and speech in the presence of multi-talker babble noise at equal intensity.

**Figure S3:**
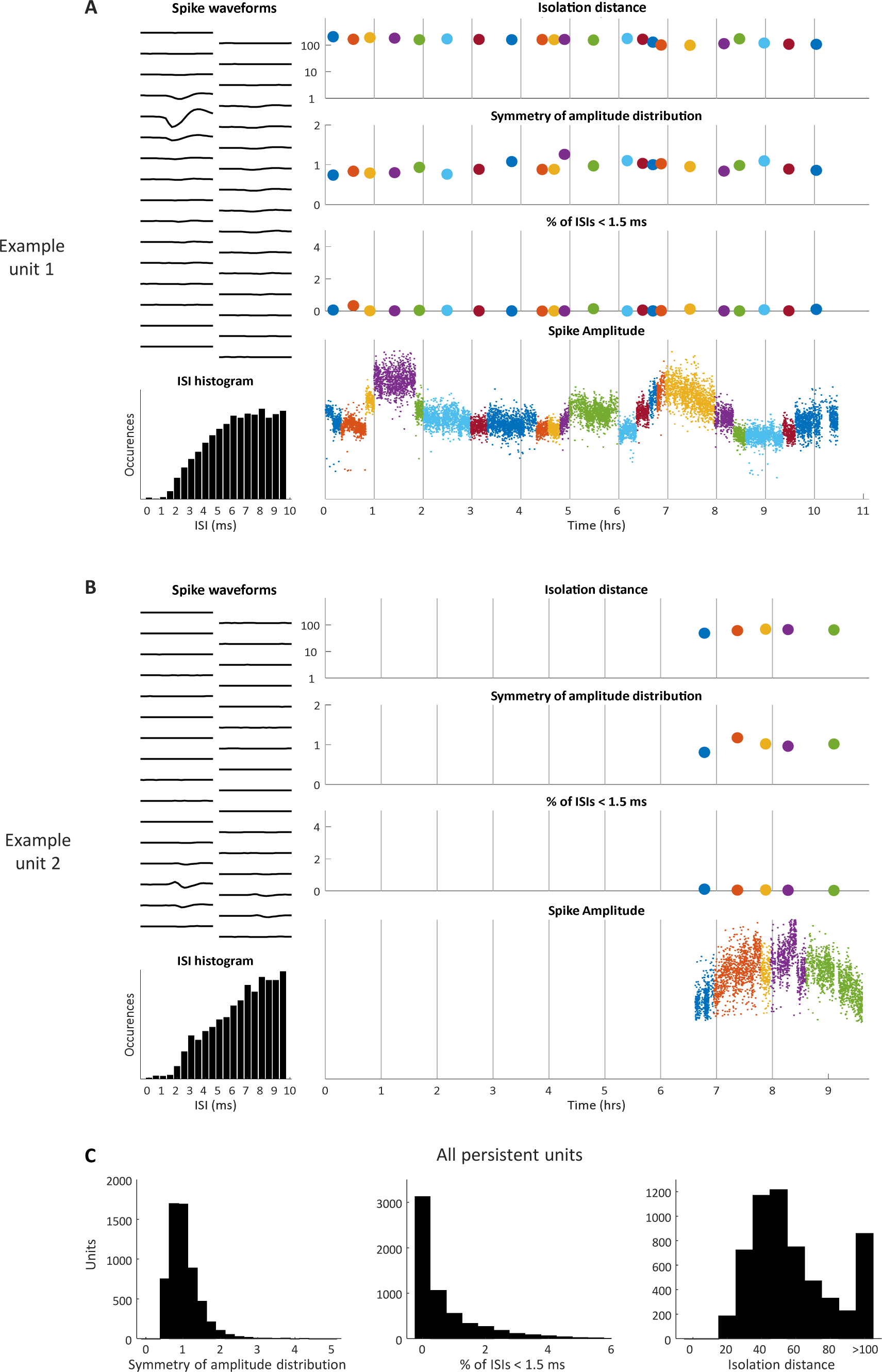
Identifying single units based on persistence. (A) An example unit that was present during an entire 10-hour recording session. The left column shows the average waveform for the unit on each of 32 electrode channels as well as the histogram of its interspike intervals. The right column shows the values of several quantities for the unit at different time points in the recording, with colors corresponding to different overlapping segments of the response as described in the Methods: (1) Isolation distance (Schmitzer-Torbert et al., 2005), which is calculated by assuming that each cluster forms a multi-dimensional Gaussian cloud in feature space and measures, in terms of the standard deviation of the original cluster, the increase in the size of the cluster required to double the number of snippets within it. A large isolation distance indicates that the cluster is well separated from other clusters, with a value of 20 typically used as a threshold for classifying a cluster as a single unit. (2) The symmetry of the spike amplitude distribution, which is measured as (a_16_ - a_2.5_)/(a_97.5_ - a_84_), where a_x_ is the spike amplitude corresponding the x^th^ percentile of the distribution of all amplitudes for that unit. A value significantly less than 1 indicates that the amplitude distribution has been truncated, i.e. that the threshold for spike detection is not low enough to capture all spikes from the unit. (3) The percentage of interspike intervals that are less than 1.5 ms, the typical absolute refractory period for IC neurons. A large value indicates that the cluster contains spikes from more than 1 unit. (4) The RMS amplitude of every spike waveform. (B) A second example unit that was only identified during the latter stages of a recording. (C) Histograms of amplitude symmetry, percentage of interspike intervals < 1.5 ms, and isolation distance for all clusters that were continuously present in a recording for at least 2.5 hours.

**Figure S4:**
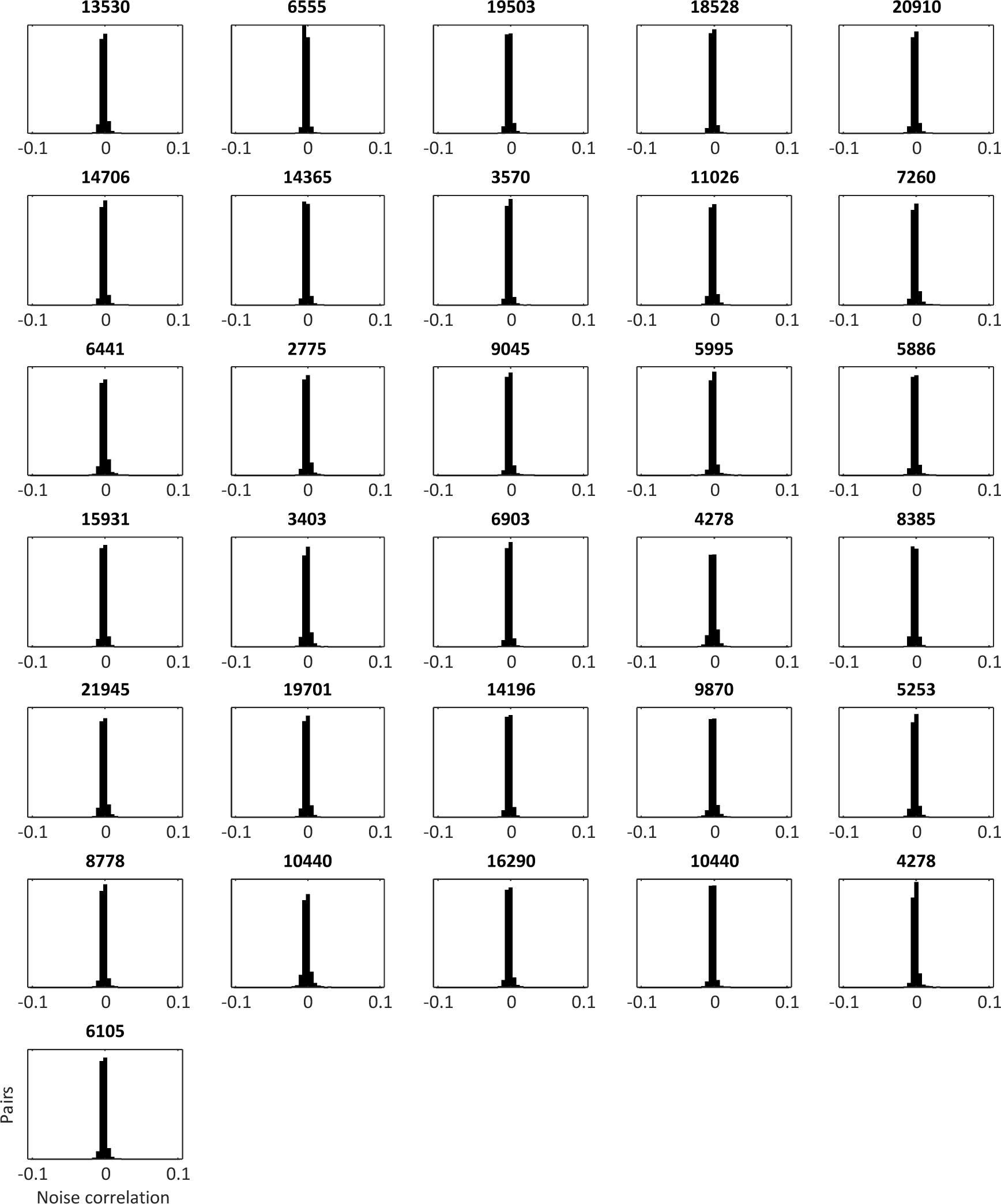
Noise correlations in IC populations are negligible. Each panel shows the distribution of noise correlations in the responses to repeated trials of identical speech at 62 dB SPL of simultaneously recorded pairs of neurons from one animal. For each neuron on each trial, the noise was measured as the difference between the response on that single trial and the mean response across repeated trials. The total number of pairs for each experiment is indicated above each panel. Only the 31 of 35 animals for which repeated trials of identical speech were presented are shown.

**TABLE S1:**
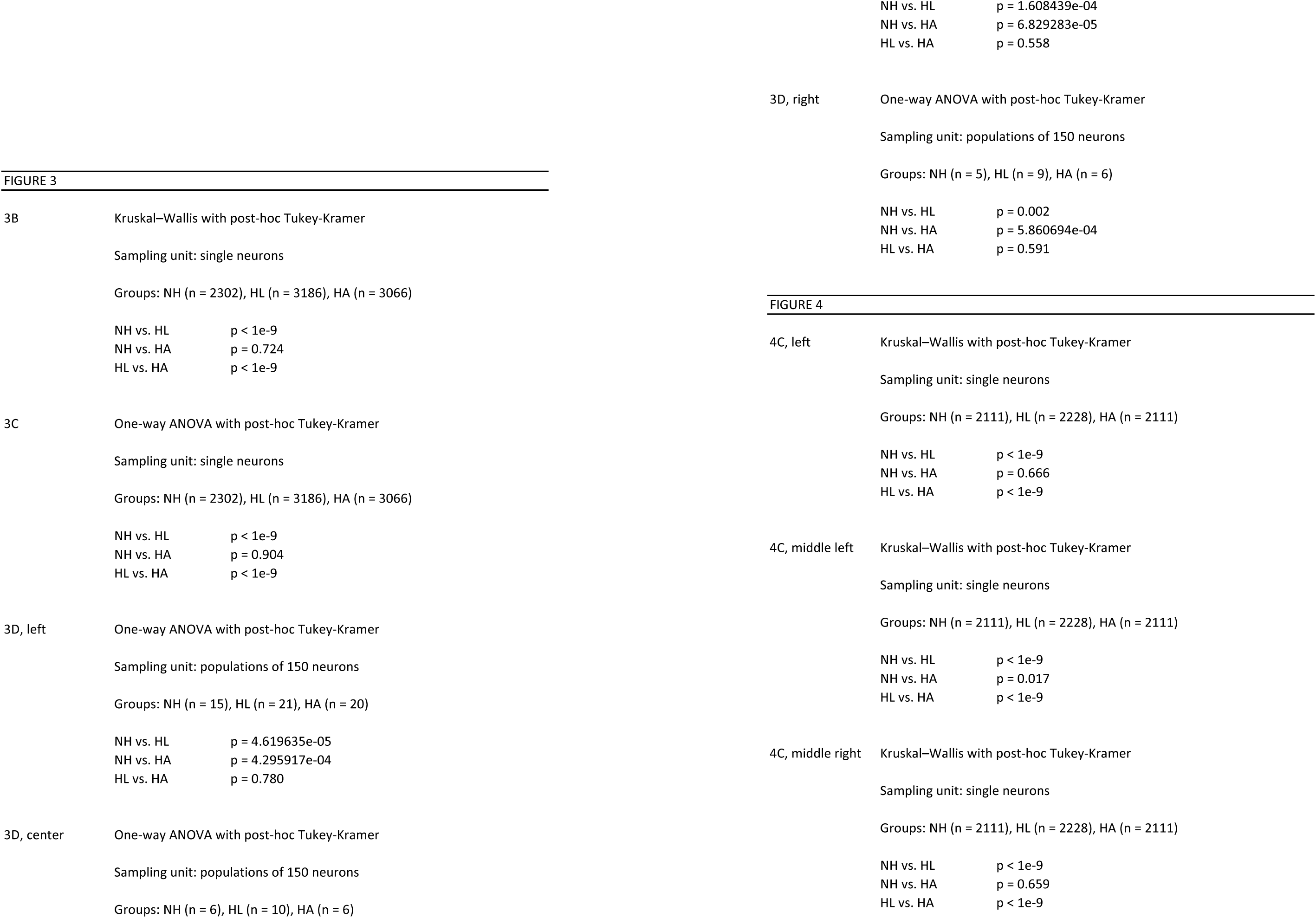
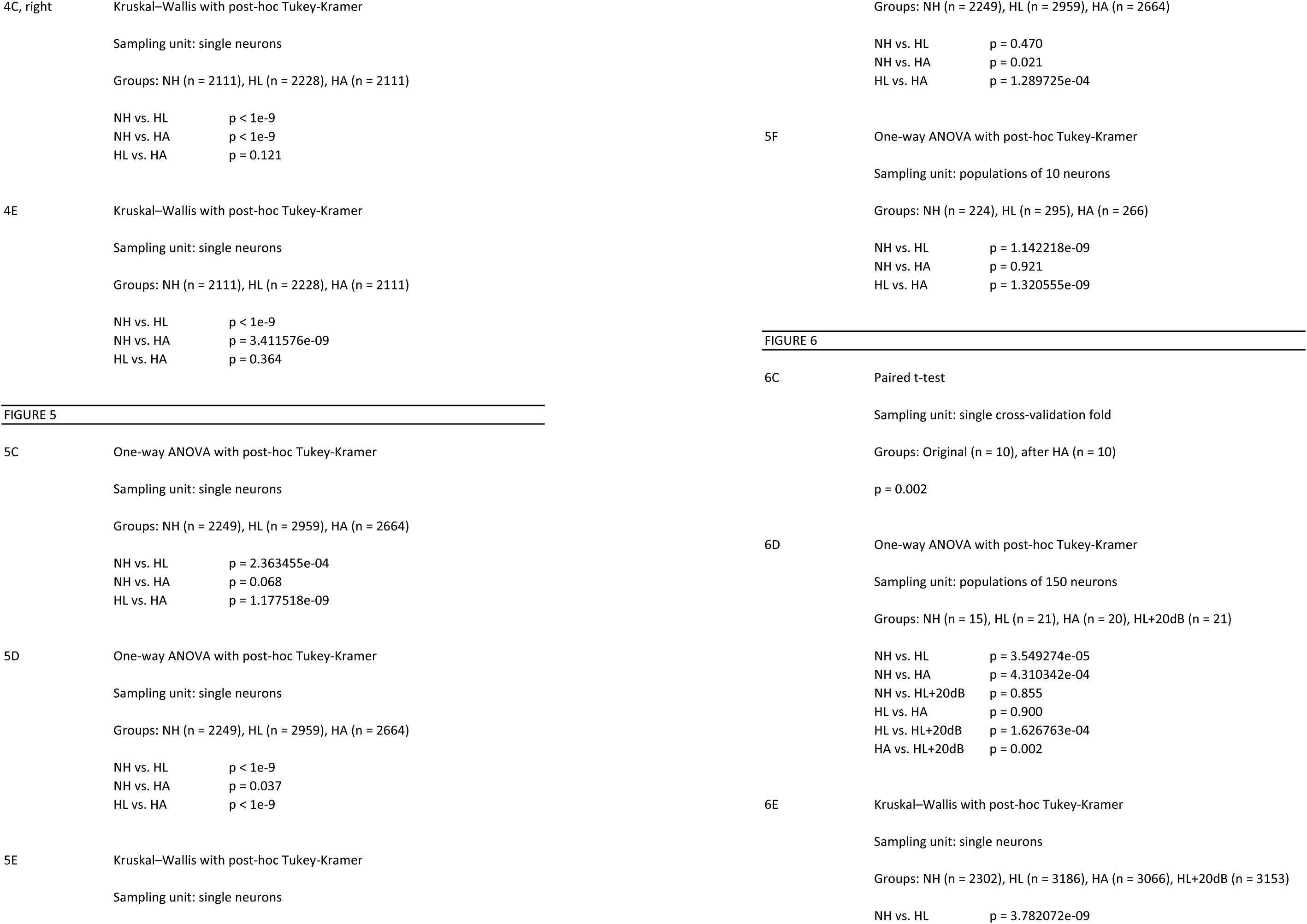
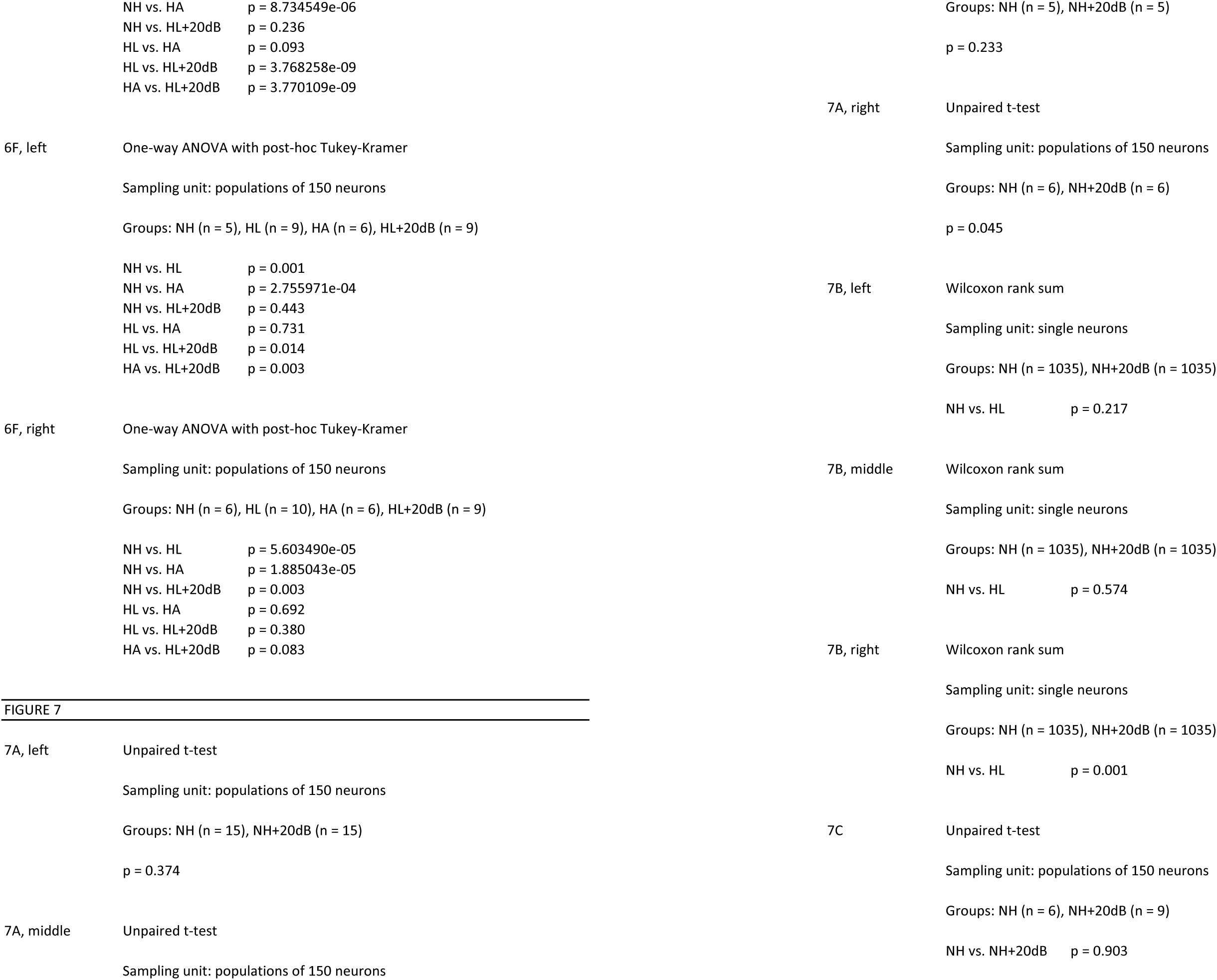
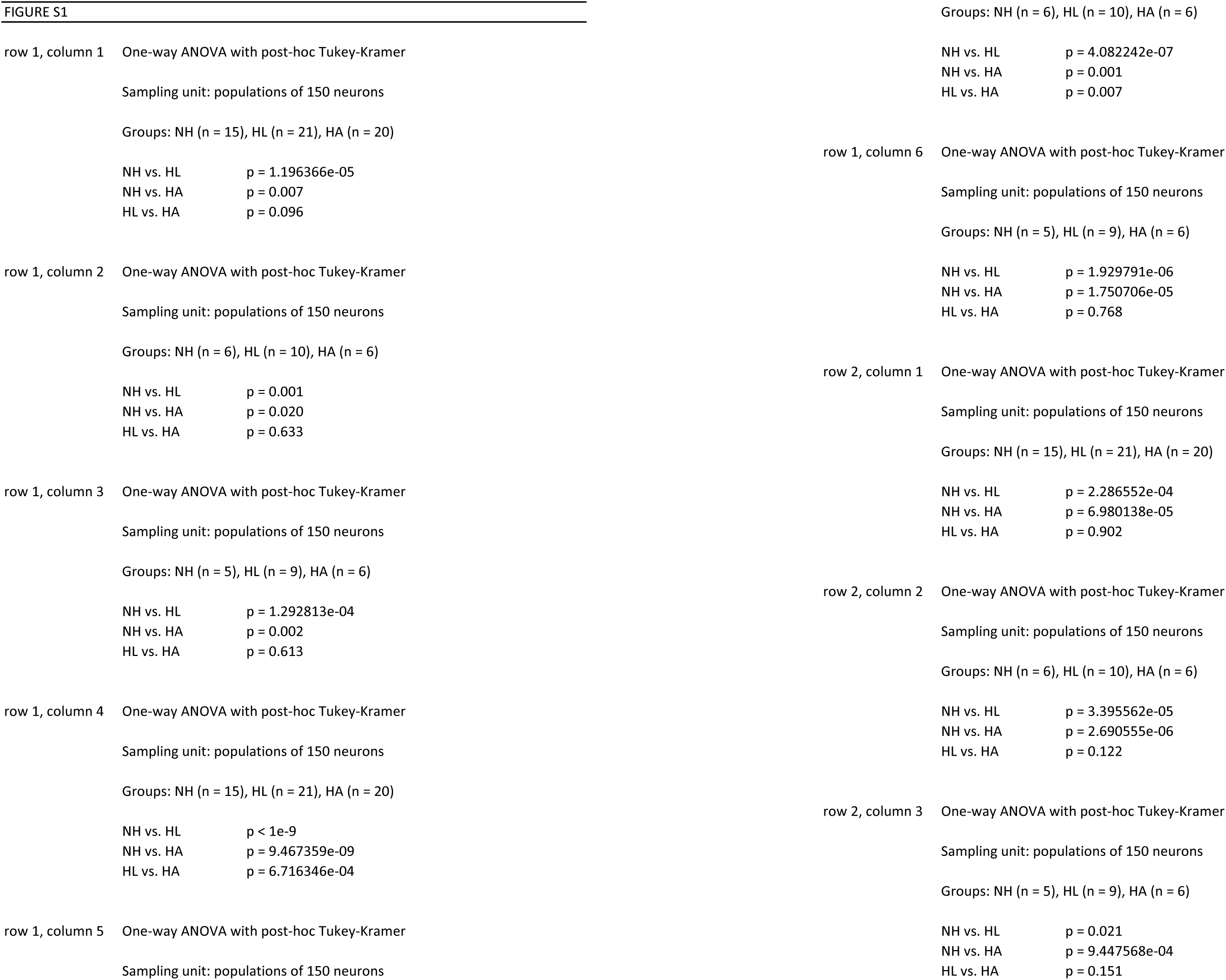
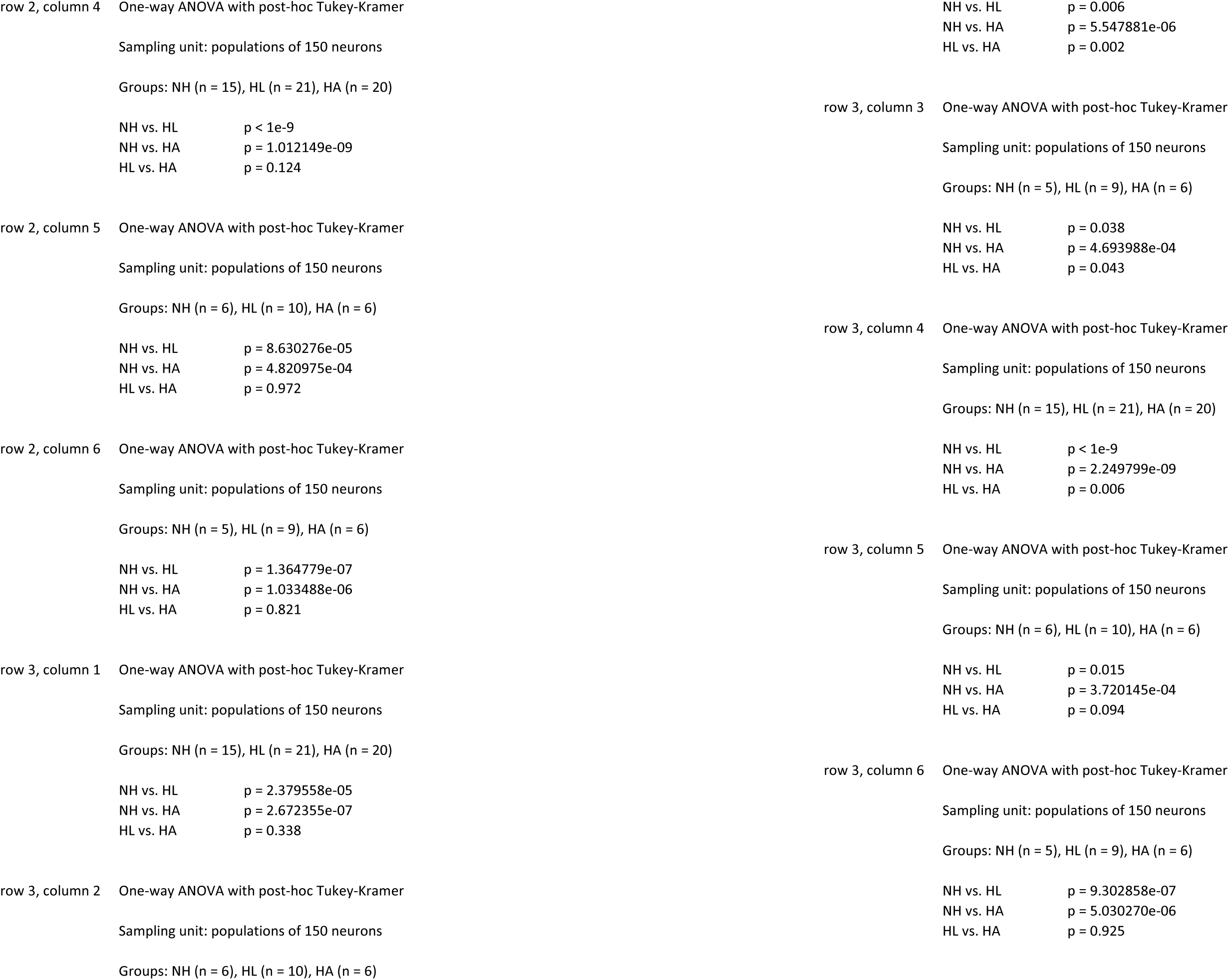
Details of statistical analyses This table provides the details of the statistical tests used in this study, including test type, sampling unit, sample sizes, and p-values. For all analyses of single neuron response properties for which distributions were not necessarily normal, non-parametric tests were used. For all analyses of classifier performance with population responses, parametric tests were used. In cases where comparisons were made across more than two groups, post hoc tests were used to compute pairwise p-values.

